# Structure and reconstitution of a hydrolase complex that releases peptidoglycan from the membrane after polymerization

**DOI:** 10.1101/2020.05.21.109470

**Authors:** Kaitlin Schaefer, Tristan W. Owens, Julia E. Page, Marina Santiago, Daniel Kahne, Suzanne Walker

**Affiliations:** Department of Microbiology, Harvard Medical School, Boston, Massachusetts 02115; Department of Chemistry and Chemical Biology, Harvard University, Cambridge, Massachusetts 02138

## Abstract

Bacteria are surrounded by a peptidoglycan cell wall that is essential for their survival^1^. During cell wall assembly, a lipid-linked disaccharide-peptide precursor called Lipid II is polymerized and crosslinked to produce mature peptidoglycan. As Lipid II is polymerized, nascent polymers remain membrane-anchored at one end and the other end becomes crosslinked to the matrix^2–4^. A longstanding question is how bacteria release newly synthesized peptidoglycan strands from the membrane to complete the synthesis of mature peptidoglycan. Here we show that a *Staphylococcus aureus* cell wall hydrolase and a membrane protein containing eight transmembrane helices form a complex that acts as a peptidoglycan release factor. The complex cleaves nascent peptidoglycan internally to produce free oligomers as well as lipid-linked oligomers that can undergo further elongation. The polytopic membrane protein, which is similar to a eukaryotic CAAX protease, controls the length of these products. A 2.6 Å resolution structure of the complex shows that the membrane protein scaffolds the hydrolase to orient its active site for cleavage of the glycan strand. We propose that this complex serves to detach newly-synthesized peptidoglycan polymer from the cell membrane to complete integration into the cell wall matrix.

---

The biosynthesis of the bacterial cell wall has been the focus of intense study for decades.^1^ The peptidoglycan precursor Lipid II is synthesized inside the cell, transported across the cytoplasmic membrane,^5,6^ and then assembled outside the cell into a crosslinked polymer that prevents osmotic lysis (Fig. 1a, left and middle panel)^2^. Two conserved families of peptidoglycan synthases carry out Lipid II polymerization and crosslinking^2–4^. These peptidoglycan synthases, particularly the transpeptidase (TP) components, have received a great deal of attention as targets for antibiotics. Indeed, transpeptidases are generically known as penicillin’binding proteins because they react covalently with beta-lactam antibiotics (pBPs)^7^. A necessary step in peptidoglycan biosynthesis that has received almost no attention is the release of newly synthesized peptidoglycan strands from the membrane (Fig. 1a, right panel). One way in which this might be achieved is with a glycosidase that cleaves within a glycan strand to separate the end that has been crosslinked into the cell wall from the lipid-linked nascent oligomer that is not yet crosslinked.^8^ Here, we describe a membrane protein complex comprising a glycosidase and a membrane protein that regulates its cleavage activity. This complex meets criteria for a peptidoglycan release factor and explains how nascent peptidoglycan is freed from the membrane in *S. aureus.*

**Figure 1.**
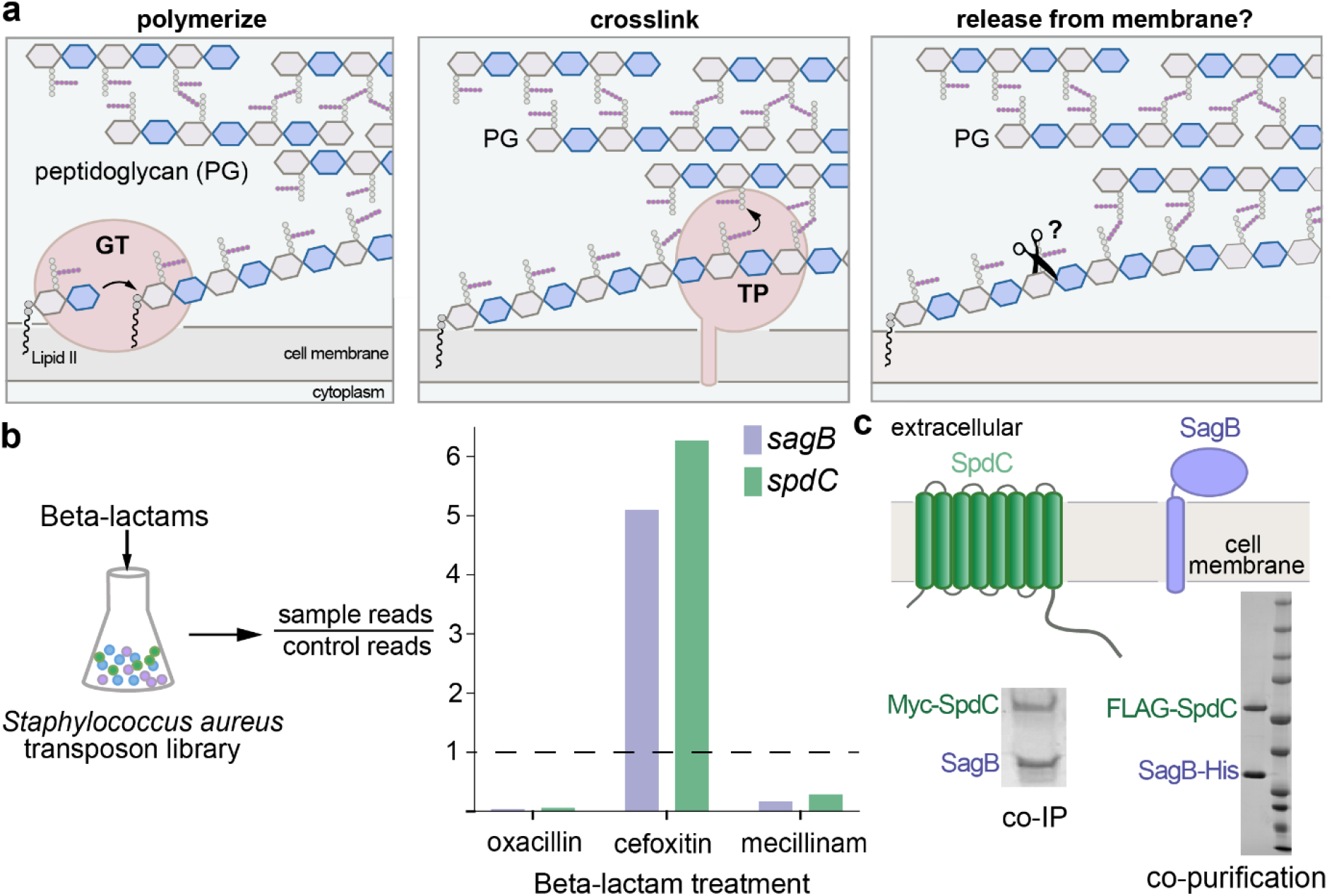
The cell wall hydrolase SagB and the membrane protein SpdC form a complex. **a,** Overview of the final steps in peptidoglycan assembly. Left panel: After translocation to the outer face of the cytoplasmic membrane, the peptidoglycan (PG) monomer Lipid II is polymerized into linear glycan strands by glycosyltransferases (GT). Middle panel: Transpeptidase domains (TPs) crosslink glycan strands into the cell wall. Right panel: The glycan strand must be released from the membrane through some type of cleavage process in order to be incorporated into the cell wall. **b,** A *Staphylococcus aureus* transposon library^9,10^ was treated with a panel of beta-lactams (oxacillin, cefoxitin, mecillinam) that have different selectivities for the four native *S. aureus* penicillin-binding proteins (PBPs)^11^. Only two genes, *sagB* and *spdC,* displayed a response pattern in which transposon reads were depleted under oxacillin and mecillinam treatment but enriched under cefoxitin treatment. **c,** SagB is a membrane-anchored glucosaminidase^12,13^ and SpdC is an eight-pass membrane protein^15^. Myc-tagged SpdC was expressed in a *△spdC S. aureus* strain and coimmunoprecipitated from solubilized membranes. (“Co-IP”, see also Supplementary Fig. 1). SagB was identified by LC-MS-MS analysis (Supplementary Table 1). Tandem affinity purification of SagB-His_6_ and FLAG-SpdC from *E. coli* yielded a stable 1:1 complex (“co-purification”, see also Supplementary Fig. 2).

To find genes important in cell wall assembly, we probed a *S. aureus* transposon library with sublethal concentrations of three different beta-lactams that have distinct PBP inhibition profiles (Fig. 1b)^9–11^. For *sagB,* encoding a membrane-anchored glucosaminidase that affects peptidoglycan strand length^12–14^, we observed an unusual response pattern in that transposon reads were strongly depleted in the presence of oxacillin and mecillinam, but enriched in the presence of cefoxitin (Fig. 1b; Supplementary Table 1). Only one other gene displayed the same pattern: *spdC,* encoding a membrane protein similar to eukaryotic CAAX proteases, enzymes that cleave prenylated proteins C-terminal to the site of prenylation. CAAX protease homologs are widespread in bacteria, but their roles have been unclear because protein prenylation is a modification not found in bacteria. Notably, *sagB* and *spdC* knockouts were reported in separate studies to share several distinctive phenotypes^12,13,15,16^. Together with the shared Tn-seq profiles, these joint phenotypes led us to think SagB and SpdC may act in a complex.

To test whether SagB and SpdC form a complex, we expressed Myc-SpdC in *S. aureus* Δ*spdC* and immunoprecipitated the tagged protein from solubilized membranes. Polyacrylamide gel electrophoresis (PAGE) of the sample showed a band that contained SagB as a major component (Fig. 1c; Supplementary Fig. 1, Supplementary Table 2). Based on this finding, we co-expressed SagB-His_6_ and FLAG-SpdC in *E. coli* and purified a complex containing SpdC and SagB in a 1:1 ratio (Fig. 1c, and also see Supplementary Fig. 2). *S. aureus* contains only one other glucosaminidase with a transmembrane helix, SagA.^12,13^ The protein is homologous to SagB but did not have the same profile in our Tn-seq experiments, and we were unable to copurify SpdC with SagA (Supplementary Fig. 3). Taken together, our experiments showed that SpdC and SagB form a stable, specific complex.

To determine whether SpdC affects SagB activity, we compared the cleavage activity of the complex and SagB alone. We incubated the enzyme or enzyme complex with uncrosslinked peptidoglycan prepared in vitro from synthetic [^14^C]-Lipid II^17–19^ and analyzed the reactions via PAGE-autoradiography (Fig. 2, and see Supplementary Fig. 4). Consistent with previous findings, SagB lacking its TM helix has low activity (Fig. 2b, lane 6; also Supplementary Fig. 5)^13,20^. In contrast, full-length SagB fully converted the peptidoglycan oligomers to diffuse bands high in the gel (Fig. 2b, lane 4). SagA produced similar product bands. These were found to be short cleavage products ranging from two to eight sugars in length (Supplementary Fig. 6). The SagB-SpdC complex also produced short oligosaccharides, but they were longer on average than those produced by SagB alone (Supplementary Fig. 6); moreover, we observed an accumulation of faster-migrating cleavage products not observed for SagB alone (Fig. 2b). We observed a similar accumulation of fast-migrating products when the native *S. aureus* substrate was used to make peptidoglycan polymer (Supplementary Fig. 7).

**Figure 2.**
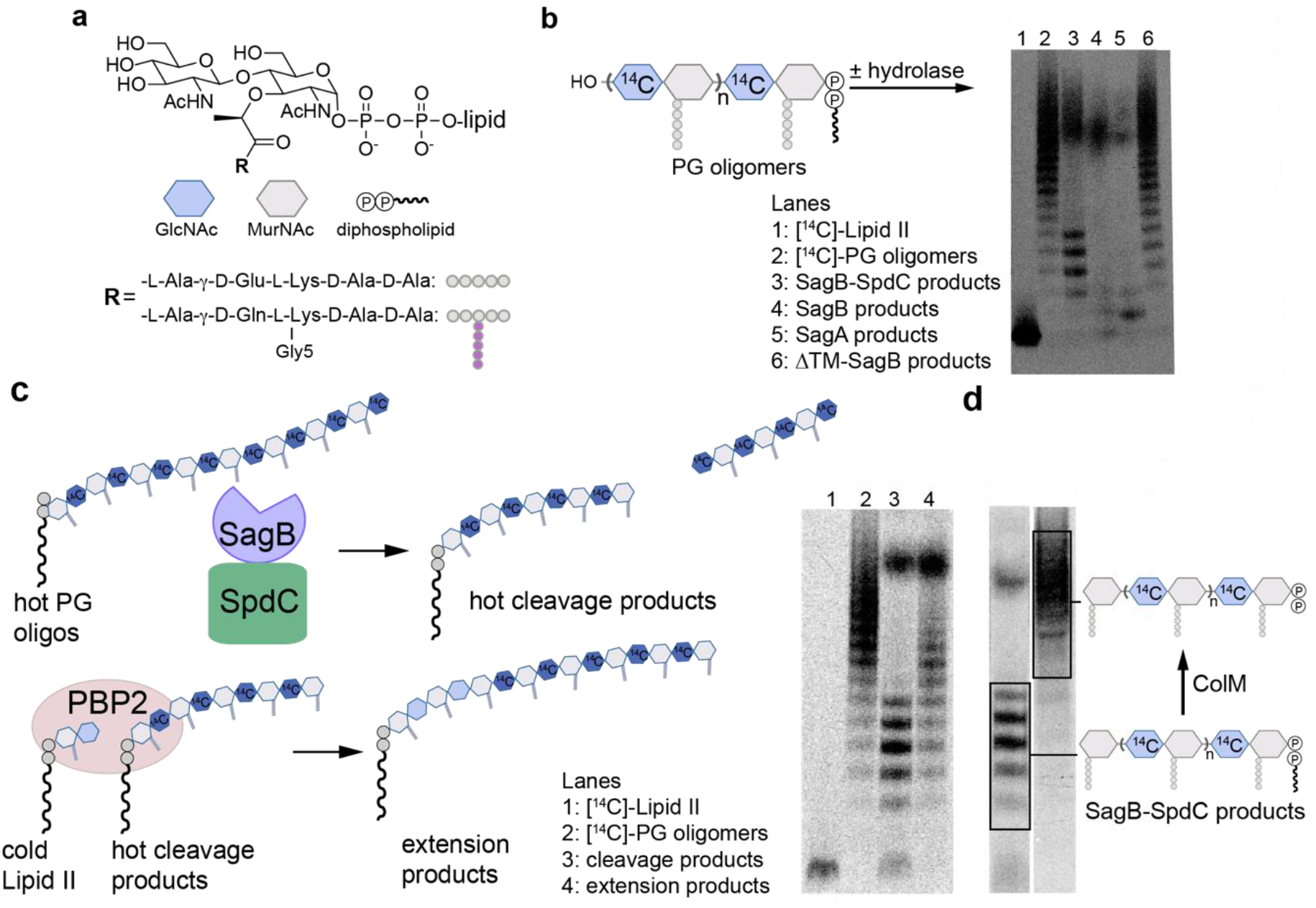
*In vitro* reconstitution shows that SagB-SpdC cleaves nascent peptidoglycan to short lipid-linked oligomers that can be elongated. **a,** Chemical and cartoon representations of the synthetic Lipid II analog^18^ and native *S. aureus* Lipid II that were used to prepare peptidoglycan polymers in panels b-d. **b,** Radiolabeled peptidoglycan polymers were incubated with the SagB-SpdC complex, SagB alone, SagA, or SagB lacking its transmembrane helix. The signal towards the top of the autoradiograph in lanes 3-5 corresponds to short, lipid-free peptidoglycan fragments, but the distribution of lengths differs (Supplementary Fig. 6). SagB-SpdC also produces a short ladder of radiolabeled peptidoglycan fragments (see also Supplementary Fig. 5). **c,** Left: schematic of assay to determine whether SagB-SpdC product bands contained a lipid-anchor. Right: Radiolabeled SagB-SpdC products were incubated with unlabeled Lipid II and PBP2 and were extended to longer products. **d,** The bacteriocin colicin M (colM) de-lipidates Lipid II and peptidoglycan oligomers, but leaves the anomeric diphosphate (Supplementary Figs. 8, 9)^23,24^. Incubation of SagB-SpdC products (lane 3) with ColM (lane 4) resulted in the complete disappearance of fast-migrating bands and the appearance of slower-migrating products. Product characterization by LC-MS confirmed the indicated structure (Supplementary Fig. 10). The faster migration of the SagB-SpdC products containing a lipid may be due to SDS binding to the lipid and increasing the net negative charge of these species.

We next sought to identify the cleavage fragments that uniquely accumulate for the SagB-SpdC complex. Their migration behavior suggested these fragments may still contain the diphospholipid anchor at the reducing end. Because peptidoglycan glycosyltransferases (GTs) add Lipid II to the reducing end of the growing polymer^21–23^, one way to test if the SagB-SpdC cleavage products retain the diphospholipid is to determine whether they are competent substrates for polymer extension (Fig. 2d). Therefore, we prepared radiolabeled peptidoglycan oligomers, cleaved them with SagB-SpdC, incubated the cleavage products with *S. aureus* PBP2 and cold Lipid II, and analyzed the products by PAGE autoradiography (Fig. 2d). The SagB-SpdC cleavage products shifted to higher molecular weight bands, showing they were competent substrates and implying the presence of a diphospholipid at the reducing end. Furthermore, when we treated the cleavage products with the bacteriocin colicin M, which removes the lipid, we observed that the cleavage products migrated more slowly by SDS-PAGE (Fig. 2e; Supplementary Fig. 8-10).^24,25^ LC-MS analysis confirmed the presence of a diphosphate on a peptidoglycan fragment having an odd number of sugars, consistent with glucosaminidase cleavage to leave a terminal MurNAc (Supplementary Fig. 10). Notably, SagB-SpdC’s cleavage activity *in vitro* depended on the conserved catalytic glutamate^13,26 13,26 13,25 13,25,20^ in SagB (E155), but not on an SpdC residue conserved in eukaryotic CAAX proteases and required for their proteolytic activity (Supplementary Fig. 11). Moreover, cellular phenotypes of the SagB catalytic mutant resemble a *sagB* knockout, whereas SpdC mutants lacking putative catalytic residues resemble wild-type (Supplementary Fig. 12)^15^. These findings show that SagB plays a catalytic role in peptidoglycan cleavage while SpdC plays a noncatalytic role in controlling SagB. While we cannot exclude the possibility that SpdC has a catalytic function, its known *in vitro* and cellular phenotypes involve noncatalytic functions.

These results show that the SagB-SpdC complex satisfies criteria expected for a peptidoglycan release factor. First, the complex yields products that contain a reducing-end lipid; second, these products are capable of further elongation. SagB’s known effect on glycan strand length may also be due to its role as part of a release factor complex. If so, we would expect the loss of SpdC to similarly affect glycan strand length. By comparing glycan strands isolated from wild-type, Δ*sagB,* and Δ*spdC* cells, we found that the short glycan strands characteristic of wild-type *S. aureus* are lost in both the Δ*sagB* and Δ*spdC* mutants (Supplementary Fig. 13), showing that SpdC is also involved in glycan strand length control in cells. We would also expect SagB-SpdC to act on newly synthesized sections of peptidoglycan polymer that are not yet modified or crosslinked into the matrix because both proteins are anchored in the membrane. Consistent with this expectation, we observed a clear preference for cleavage of nascent, unmodified peptidoglycan over peptidoglycan that contained teichoic acid modifications or crosslinks (Supplementary Fig. 14 and 15). Our conclusion that SagB-SpdC acts early in the peptidoglycan maturation pathway is supported by recent studies that used atomic force microscopy to visualize SagB mutants^14^.

To understand how SpdC interacts with SagB to control cleavage of nascent polymer, we solved the structure of SagB complexed with a truncated form of SpdC (SpdC^1-256^) that lacks the cytoplasmic C-terminal domain (Fig. 3a-c; Supplementary Fig. 17)^15^. Removal of this apparently unstructured domain did not affect formation of the complex or change its *in vitro* activity, nor did it impact cellular phenotypes of SpdC (Supplementary Fig. 11, 16, 17). We refer hereafter to this truncated complex simply as SagB-SpdC. Our structure, obtained using lipidic cubic phase (LCP) crystallography^27^, resolves nearly all of SpdC, which contains 8 transmembrane helices linked by short extracellular and cytoplasmic loops, as well as the transmembrane helix and glucosaminidase domain of SagB (Fig. 3c).

**Figure 3.**
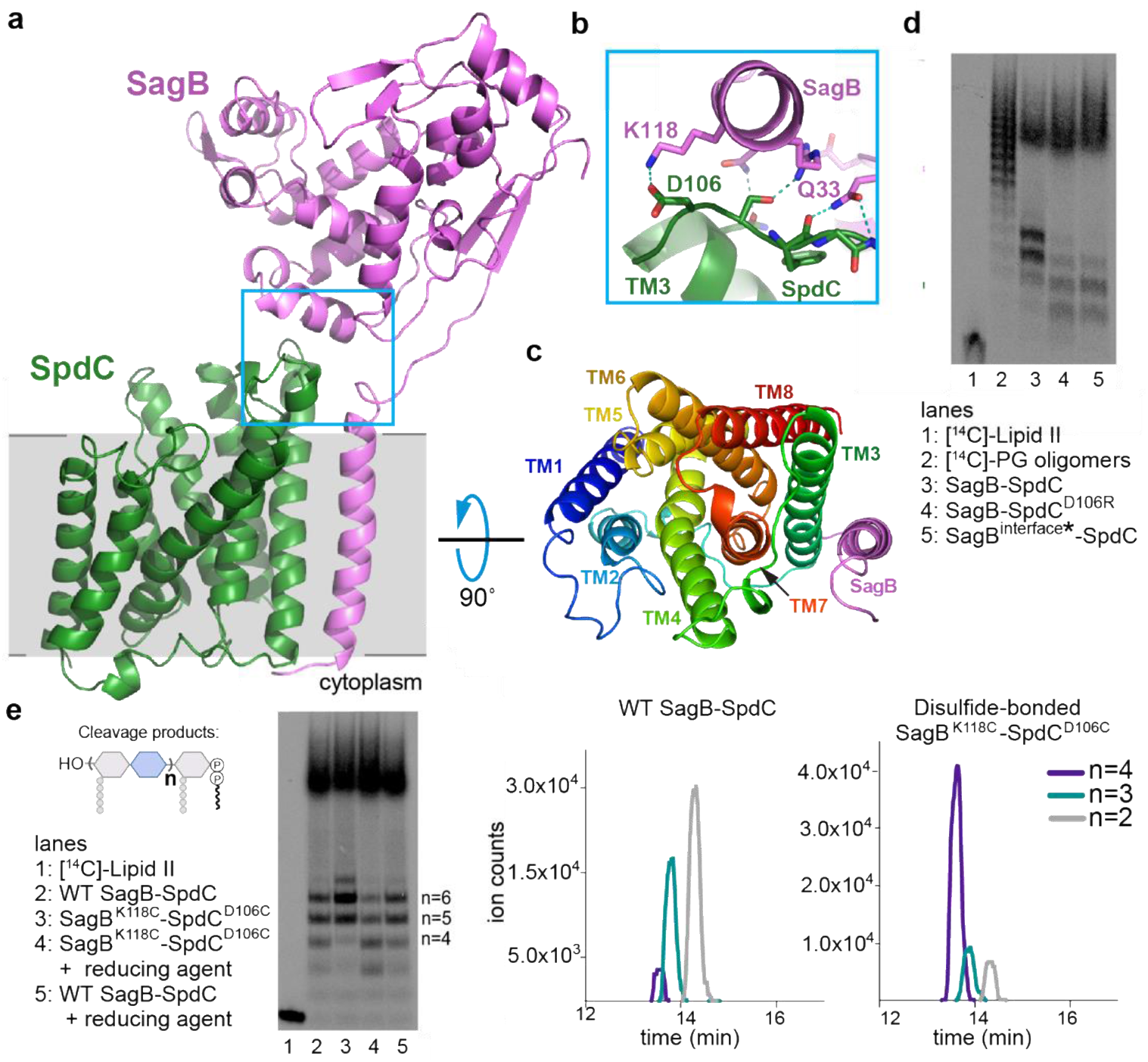
A 2.6 Å resolution crystal structure of the SagB-SpdC complex establishes that two interfaces are critical for its function. **a,** A cartoon representation of the SagB-SpdC crystal structure. The extracellular domains of both proteins interact (blue box); a helix at the bottom of the active site cleft of SagB (violet) contacts an extracellular loop between TM3 and TM4 of SpdC (green). The approximate location of the membrane is denoted in gray. **b,** Several hydrogen bonds and a salt-bridge form at the interface between the SagB helix and the SpdC loop. **c,** A view from the extracellular face of the transmembrane helices shows that SagB closely contacts TM3 of SpdC. SagB lacking its TM helix does not co-purify with SpdC (Supplementary Fig. 19). **d and e,** Radiolabeled peptidoglycan oligomers were incubated with SagB-SpdC or with constructs containing mutations designed to either disrupt or stabilize the extracellular interface between SagB and SpdC. **d,** SagB^interface^* denotes SagB^N115SA, K118N, R119Q, V122D, D123G, L127E^, in which SagB residues at the interface were switched to the corresponding SagA residues. **e,** A variant of SagB-SpdC with two cysteine-substituted residues, SagB^K118C^ - SpdC^D106C^, was purified as the disulfide-linked complex (Supplementary Fig 20). Activity of the oxidized complex (lane 3) was compared to the activity of SagB^K118C^-SpdC^D106C^ incubated with reducing agent (lane 4), and to wild-type SagB-SpdC without and with reducing agent (lanes 2 and 5). Unlabeled cleavage products were also treated with ColM and analyzed using LC-MS analysis. Extracted ion chromatogram (EIC) traces are shown for both wild-type SagB-SpdC and SagB^K118C^-SpdC^D106C^ reactions, and further confirm that longer oligosaccharide products are preferred for a disulfide-restricted complex. Notably, short oligosaccharides ionize better relative to longer oligosaccharides.

SagB makes contacts to SpdC on both its extracellular face and in the membrane (Fig. 3a-c). A helix and loop in SagB’s glucosaminidase domain (residues 115 to 127) sit over the edge of the SpdC helical bundle and make extensive hydrogen bonding and ionic interactions with the surface-exposed loop between SpdC transmembrane helix 3 (TM3) and TM4 (Fig. 3b). Adjacent to this interface, the top of the SagB transmembrane helix also contacts the SpdC TM3-TM4 loop (Q33, Fig. 3b), and from there the SagB TM helix maintains tight hydrophobic contacts with SpdC TM3 across the membrane (Fig. 3c, and also see Supplementary Fig. 19). We found that the SagB TM helix is required to form the complex: SpdC did not co-purify with soluble SagB and swapping the TM helix of SagB with that of SagA greatly reduced stability of the complex (Supplementary Fig. 19). However, the SagB TM helix is not sufficient for robust complexation with SpdC; when we replaced the TM helix of SagA with that of SagB, they did not co-purify as a 1:1 complex (Supplementary Fig. 19). Taken together, our results show that the transmembrane interactions are necessary to form a complex, but are not sufficient for the activity displayed by SagB-SpdC.

We thought the contacts at the extracellular SagB-SpdC interface could affect cleavage function and made mutations predicted to disrupt key interactions. SagA and SagB share the same fold and are 53% similar (Supplementary Fig. 3), but the extracellular residues in SagB that contact SpdC are not conserved in SagA. Replacing these residues on the extracellular SagB helix that contacts SpdC with the corresponding SagA residues (SagB^115-SEVNQLLKG-123^) did not prevent complex formation, which may be driven largely by the interactions between TM helices in the membrane; however, we observed an erosion of product length control (Fig. 3d, lane 5). We identified a possible salt bridge between SagB lysine 118 and SpdC aspartate 106 in the crystal structure (Fig. 3b), and found that replacing either of these amino acids with a residue having the opposite charge also resulted in a product distribution more closely resembling that of purified SagB alone (Fig. 3d, lane 4). These results suggested that the relative orientation of SpdC and the SagB glucosaminidase domain is important for determining the product distribution.

To test whether the interface conformation observed in the crystal structure is critical for product length control, we generated disulfide-linked SagB-SpdC complexes using the crystal structure as a guide. We mutated proximal interface residues in SagB and SpdC to cysteines and purified the corresponding complexes (Supplementary Fig. 20a). Both SagB^N115C^-SpdC^S107C^ and SagB^K118C^-SpdC^D106C^ complexes formed disulfide linkages as judged by SDS-PAGE analysis. Both disulfide-linked complexes produced an altered distribution of lipid-linked peptidoglycan products compared to wild-type (Fig. 3e and Supplementary Fig. 20), with a shift to longer products. By comparing the mobility of the lipid-linked SagB-SpdC cleavage products to Lipid II and short oligomers, we concluded that the cleavage products from the disulfide-bonded complexes have 9-13 sugars (n=4-6; Supplementary Fig. 20). In an analogous experiment, unlabeled peptidoglycan oligomers were treated with wild-type or the disulfide-bonded complex, followed by ColM treatment and then LC-MS analysis. Similar to PAGE autoradiography, the predominant species were longer for the reaction with the disulfide-bonded complex (Fig. 3e). These results show that restricting the orientation of the two proteins to that present in the crystal structure results in tighter length control. Noting that the crystal structure was obtained in a membrane-like environment, we infer that this orientation is relevant to the mechanism of cleavage in cells.

Our results suggest a mechanism for how SagB-SpdC generates the observed product lengths (Fig. 4). The structure of SpdC is similar to that of the CAAX protease Rce1 (Supplementary Fig. 21), which cleaves prenylated proteins just after the modified cysteine residue. The resemblance of SpdC to a CAAX protease suggests that SpdC may bind part of the lipid pyrophosphate carrier of nascent peptidoglycan. SpdC contains a cavity that opens to the membrane between TM1 and TM5, a possible site of prenyl chain entry. A groove extends from this opening along the extracellular face of SpdC all the way to the active site pocket of SagB. Several lines of evidence have established the directionality of oligosaccharide binding for the family of glucosaminidases to which SagB belongs^28^, and this directionality is consistent with nascent peptidoglycan following the groove such that the non-reducing end of the polymer exits toward the cell wall (Supplementary Fig. 22). Consistent with our model, we found that the distribution of nascent cleavage products was unchanged whether polymerase and SagB-SpdC were added simultaneously or sequentially (Supplementary Fig. 23). These results imply that cleavage occurred after polymer release from the polymerase, leaving the substrate lipid tail accessible for SpdC to bind. The product lengths observed *in vitro* are in good agreement with the lengths that would be predicted from the physical dimensions of the complex if the polymer tracks along the grooves in SpdC and SagB (Fig. 4b). To determine if SagB-SpdC activity depends on the presence of the lipid portion of oligomers, we pre-treated peptidoglycan with ColM and incubated the mixture with SagB-SpdC. As analyzed by PAGE autoradiography, SagB-SpdC activity is abrogated when oligomers lack the lipid, consistent with a role for the lipid in interacting with the complex (Supplementary Fig. 24). Obtaining a structure of SagB-SpdC bound to a lipid-linked nascent oligomer is now a key goal.

**Figure 4.**
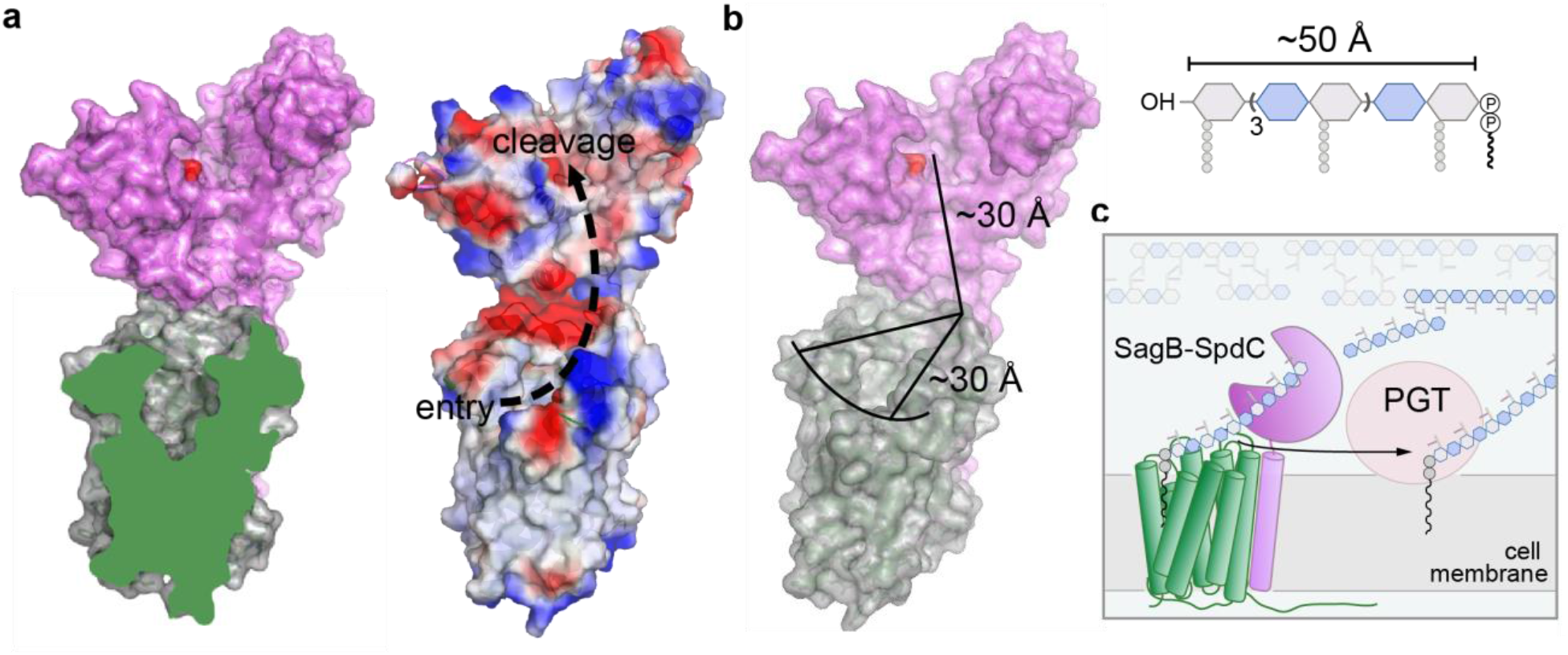
SagB-SpdC is a peptidoglycan release factor that cleaves at a defined length from the reducing end to allow strands to be fully incorporated into the cell wall. **a,** A cross-section of SpdC shows a groove that extends from the membrane and into the SagB catalytic groove (also see Supplementary Fig. 21d). As depicted by electrostatic surface potential, SpdC provides a path for a glycan strand that extends from an “entry” point in the membrane and into the “cleavage” site as denoted by the catalytic glutamate (red). **b**, Measuring the distance along this path provides approximately equivalent lengths required for binding a glycan strand that represents the average lipid-linked product lengths observed for the SagB-SpdC complex. **c**, Scheme depicting the proposed mode by which SagB-SpdC could act as peptidoglycan release factor. To prevent wasteful release of lipid-free peptidoglycan fragments, SagB-SpdC cleaves short strands that are being or have been crosslinked into the cell wall matrix; lipid-linked peptidoglycan products can be further elongated by peptidoglycan polymerases (PGTs).

SagB-SpdC is the first example of a release factor complex shown to cleave nascent peptidoglycan from its membrane anchor, which would allow its full integration into the cell wall. As *S. aureus* can survive without SagB-SpdC, we infer that other peptidoglycan release factors exist, and some may also be membrane-anchored cell wall hydrolase complexes. We note that this first structure and biochemical analysis of a bacterial member of the CAAX protease family suggests that other CAAX protease homologs, which are widespread in bacteria, may also act to scaffold membrane proteins involved in cell envelope synthesis.

## Acknowledgements

We thank Dr. Samir Moussa for his preliminary experiments investigating the roles of SpdC and SagB. We also thank Dr. Andrew Kruse for helpful discussions on crystallography. This work used NE-CAT beamlines (GM103403), a Pilatus detector (RR029205), and an Eiger detector (OD021527) at the APS (DE-AC02-06CH11357). This research was supported by GM076710 and U19 AI109764 to D.K. and S.W. and T32GM007753 to J.E.P.

## Data availability

Transposon sequencing data (BioProject accession number PRJNA573479) can be found in the NCBI BioProject database.

## Author contributions

S.W., K.S., T.W.O., and J.E.P. designed experiments and analyzed the data with input from D.K. K.S. performed the biochemical experiments; K.S. and T.W.O. purified proteins and performed crystallographic experiments; J.E.P. analyzed transposon sequencing data, constructed *S. aureus* mutant strains, performed the spot dilutions, and performed the glycan strand experiment; S.W., K.S., T.W.O., J.E.P., and D.K. wrote the manuscript with input from all authors.

## Competing interests

The authors declare no competing interests.

## Methods and Materials

### Materials

All reagents and chemicals were purchased from Sigma-Aldrich unless indicated otherwise. Lysostaphin was purchased from Ambicin. *Staphylococcus aureus* was grown in tryptic soy broth (TSB) with aeration or on TSB with 1.5% agar at 30 or 37°C. Antibiotics were used at the following concentrations for *S. aureus* strains: kanamycin (50 μg/mL), neomycin (50 μg/mL), tetracycline (3 μg/mL), chloramphenicol (10 μg/mL), and erythromycin (10 μg/mL). NovaBlue (DE3) *Escherichia coli* (E. coli) strains were grown in Lysogeny broth (LB). BL21(DE3) *E. coli* strains were grown in terrific broth at temperatures between 18°C and 37°C as described below. Native Lipid II was extracted from *Staphylococcus aureus* as previously described^19^. Synthetic Lipid II was prepared as previously described^18^. The radiolabeled wall teichoic acid (WTA) precursor, [^14^C]-LII_A_^WTA^, was prepared as previously reported^17^. FMOC-Biotin-D-Lysine (BDL) was used to prepare BDL^29^. PBP2^S398G^, SgtB^Y181D^, *Ef*PBPX, and TagT were expressed and purified as described in previous methods^19,30–32^. Colicin M was expressed and purified as previously described^5^. LytA was expressed and purified as previously described^33^.

## Methods

### Beta-lactam treatment and sequencing of a methicillin-resistant *Staphylococcus aureus* transposon library

We created a high-density transposon library in *S. aureus* USA300 by phage-based transposition as previously described^9,10,34^. The transposon library was treated with beta-lactams (8 μg mL^-1^ for mecillinam, 0.4 μg mL^-1^ for cefoxitin, and 0.1 μg mL^-1^ for oxacillin) at 37°C. A low concentration of oxacillin was chosen to identify factors important for beta lactam resistance. For mecillinam and cefoxitin, the concentrations were chosen such that only one PBP should be significantly inhibited in each condition. For mecillinam, the concentration was that at which the treated cells phenocopied a *pbp3* mutant at 43 °C. For cefoxitin, the concentration was that which sensitized cells to oxacillin to the same degree as a *pbp4* mutant. Cultures were shaken until A280=1-2.0. Cells were spun down, and DNA was isolated in preparation for Tn-Seq as previously described. Using reported methods^9,10^, genes significantly enriched and depleted under cefoxitin, mecillinam, or oxacillin conditions were identified using a two-sided Mann-Whitney U test corrected for multiple hypothesis testing using the Benjamini-Hochberg method. A gene was considered to be enriched if the treated:control read ratio was greater than five and depleted if the treated:control read ratio was less than 0.1. For mecillinam, the cut-off for depletion was loosened to 0.3, as mecillinam was moderately selective. Scripts for this analysis can be found at https://github.com/SuzanneWalkerLab/TnSeqMOAPrediction.

### Plasmid construction for *S. aureus* strains

#### pKFC_spdC_kan

The kan^R^ marker (primers SM45 and SM46) and the 1-kb sequences upstream (primers SM43 and SM44) and downstream (primes SM47 and SM48) of the *spdC* open reading frame were amplified by PCR and stitched together by overlap PCR. The resulting fragment was cloned between the BamHI and SalI restriction sites of pKFC.^35^ *pKFC_spdC*

The 700-bp sequences upstream (primers SM1 and SM2) and downstream (primers SM3 and SM4) of the *spdC* open reading frame were amplified by PCR and stitched together by overlap PCR. The resulting fragment was cloned between the BamHI and SalI restriction sites of pKFC.

#### pJP47

The pTarKO vector was linearized with primers F_pKTarO and R_pKTarO using the plasmid pTD47^36^ as a template. The 1-kb sequences upstream (primers F_1kb+_sagB and R_lkb+_sagB) and downstream (F_1kb(-)_sagB and R_1kb(-)_sagB) of the *sagB* open reading frame were amplified by PCR. Overlap PCR was performed to assemble these fragments with the tet^R^ marker, and then the resulting fragment was ligated into the plasmid backbone between the restriction sites BamHI and SalI. Then, the tet^R^ marker (primers oJP51 and oJP52) and upper homology arm (primers oJP54 and oJP33) were sequentially replaced by Takara Bio In-Fusion seamless cloning after linearizing the plasmid with primers oJP49 and oJP50 and oJP53 and oTD145 respectively. The insert containing the *sagB* homology arms and tet^R^ marker was amplified with primers oJP32 and oJP35 and cloned between the BamHI and SalI restriction sites in pKFC. To exchange the tet^R^ marker for a kan^R^ marker, this plasmid was linearized with primers oJP79 and oJP80. The linearized DNA was phosphorylated at the 5’ ends using T4 polynucleotide kinase, and the ends were ligated to produce circular DNA. The kan^R^ marker (primers oTD73 and oTD74) was then cloned into the plasmid at the XbaI restriction site. Finally the insert containing the *sagB* homology arms and the kan^R^ marker was amplified with primers oJP32 and oJP35 and cut into the pTarKO backbone between the BamHI and SalI restriction sites^37^.

#### pSM_spdC_myc, pJP17, and pJP42

For pSM_spdC_myc and pJP42, the full or truncated *spdC* gene sequence with its native ribosome-binding site (−17) was amplified from HG003 *S. aureus* genomic DNA and an amino-terminal cMyc tag appended by PCR using primers SM165 and SM166 for pSM_spdC_myc and primers oJP25 and oJP81 for pJP42. The fragments were then cloned between the KpnI and BlpI restriction sites of pTP63.^38^ For pJP17, the cMyc-*spdC* fragment with the native ribosome binding site was amplified from pSM_spdC_myc using primers oJP25 and oJP26 and cloned between the KpnI and BlpI restriction sites of pTP63.

#### pSM_spdC_his

The *spdC* gene and native ribosome-binding site was amplified from HG003 *S. aureus* genomic DNA and a carboxy-terminal hexa-histidine tag was appended by PCR with primers SM124 and SM125. This fragment was cloned between the KpnI and BlpI restriction sites of pTP63.

#### pSM_spdC_E135A, pSM_spdC_R139A, pSM_spdC_H210A

These three plasmids were constructed using QuikChange site-directed mutagenesis with primers SM130 and SM131, SM132 and SM133, and SM134 and SM135 respectively and *pSM_spdC_his* as a template.

#### pJP15 and pJP19

The *sagB* or *sagB E155A* gene sequence with a carboxy-terminal hexa-histidine tag was amplified from *pspdC_sagB* or *pspdC_sagB^E155A^* respectively and the native ribosome binding site appended by PCR with primers oJP21 and oJP22. The fragments were cloned between the KpnI and BlpI restriction sites in pTP63.

#### pJP22

A gBlock gene fragment was synthesized by IDT. The fragment was amplified with primers oJP30 and oJP31 and cloned between the KpnI and BlpI restriction sites in pTP63.

### *S. aureus* strain construction

pKFC_*spdC_*kan and pKFC_*spdC* were used to construct SHM056 and SHM002 respectively using a previously published method^35^. pJP47 was used to construct JP132 using a previously published method^37^. The deletions, or in the case of SHM002 the integrated plasmid before recombination, were transduced to HG003 *S. aureus.* The final deletions were confirmed by colony PCR and sequencing. Phage transductions were performed using a previously published protocol^39^.

To construct JP012 and JP065, a phage lysate was prepared from SAUSA300 JE2 *sagB::*Tn-erm^R^ from the Nebraska library and used to transduce HG003 *S. aureus* and SHM056 respectively.

To construct strains containing pTP63^38^ constructs, the plasmids were first electroporated into TD011, and transformants were selected on 10 μg/mL chloramphenicol at 30°C. The pTP63 constructs were transduced from these transformants into strain JP012 to produce strains JP051 and JP053 and into SHM056 to produce SHM226, JP054, JP064, and JP128. For JP061, JP062, and JP063, the pTP63 constructs were first transduced into SHM002, and from there transduced into SHM056.

### Co-immunoprecipitation with Myc-tagged SpdC in *Staphylococcus aureus*

This protocol was adapted from previously published protocols^40^. An overnight culture of SHM226 was diluted 1:100 into 1 L of TSB. The culture was grown at 37°C with shaking at 200 r.p.m. until A_600nm_ = 0.6 and then 0.2 μM anhydrotetracycline was added to induce plasmid expression. After a 3 h induction, cells were pelleted at 5000xg, 15 minutes, 4°C. Cell pellets were then resuspended in lysis buffer (1X PBS (pH 7.4), 20 μg mL^-1^ DNase and RNase, 10 μg mL^-1^ lysostaphin, 5 mM MgCl_2_) and incubated at 37°C for 1 h. In samples treated with a chemical crosslinker, 0.5 mM DSP was added to the mixture for 1 h and then quenched with 20 mM Tris (pH 7.5). After cooling on ice, cells were lysed with a French press two times at 20,000 psi on a high ration setting. Unbroken cells were then removed by centrifugation at 10,000xg, 4°C, 15 min. Membranes were pelleted by ultracentrifugation at 100,000xg, 4°C for 60 minutes in a Beckman 45Ti rotor. For solubilization, membrane pellets were resuspended in buffer B (1X PBS (7.4), 500 mM NaCl, 1% Triton X-100). Cell membranes were then rocked at 4°C overnight before insoluble cell debris was removed by ultracentrifugation at 100,000xg, 4°C, 30 minutes. Equilibrated magnetic anti-Myc beads (Clontech, Catalog #635699) were then added to the solubilized membranes and rocked at 4°C overnight. Equilibrated beads were then washed three times with wash buffer (1X PBS pH 7.4, 200 mM NaCl, 1% Triton X-100). Protein was eluted with elution buffer provided in the Myc Immunoprecipitation kit (Clontech, Catalog # 635698). Elution buffer was then neutralized with 1 N NaOH before running on a 4-20% SDS-PAGE gel. Protein bands were prepared for LC-MS-MS analysis, adapted from previous protocols. Protein bands were excised from the gel and stored in deionized H_2_O prior to submission for LC-MS-MS analysis at the Taplin Mass Spectrometry Facility, Harvard Medical School.

### Spot dilution assay

Overnight cultures were diluted 1:100 into 3 mL TSB and grown at 30 °C with aeration until mid-log phase. Cultures were then diluted to OD_600_ = 0.5. Five 10-fold serial dilutions of the resulting cultures were prepared for each strain, and 5 μL of each dilution was spotted on TSA plates with or without 0.4 μM anhydrotetracycline inducer and, where indicated, 0.8 μg/mL tunicamycin or 1 μg/mL lysostaphin. Plates were imaged after approximately 16 hours of incubation at 30 °C. Strains HG003 wild-type, SHM056, JP012, JP051, JP053, JP054, JP061, JP062, JP063, JP064, JP065, and JP128 were used for these assays.

## Cloning, expression, and purification of *S. aureus* glucosaminidases and SpdC variants

### Cloning of *S. aureus* glucosaminidases and SpdC

Genes encoding SagA (SAV2307), SagB (SAOUHSC_01895), and SpdC (SAOUHSC_02611) were amplified by PCR from *Staphylococcus aureus* strain NCTC 8325 genomic DNA. For co-expression, SagB and SpdC were cloned into a pDUET containing an amino-terminal SUMO-fusion followed by a Flag epitope tag and a carboxyterminal hexa-histidine tag in another site. SagB and SpdC were amplified using F_SagB/R_SagB and F_SpdC/R_SpdC, and ligated into the pDUET, using primers F_DUET_SagB/R_DUET_SagB and F_DUET_FLAG_SpdC/R_DUET_FLAG_SpdC in two steps using Gibson assembly (New England Biolabs, # E2611L). Similar methods were used for SagA and SpdC co-purification. Oligonucleotide primers were purchased from Eton Bio.

### Expression and co-purification of full-length, wild-type *S. aureus* SagB and SpdC

For co-expression of *S. aureus* SagB and SpdC, overnight cultures of BL21(DE3) *E. coli* containing pDUET-SUMO-FLAG-SpdC and SagB-His_6_ and an arabinose-inducible Ulp1 protease plasmid^3^ (pAM174) were diluted 1:100 into terrific broth supplemented with 50 μg ml^-1^ carbenicillin and 35 μg ml^-1^ chloramphenicol. Cultures were grown at 37°C with shaking at 200 r.p.m. until A_600nm_ = 0.6 and then shifted to 18°C. At an A_600nm_ = 1.0, protein expression was induced by addition of 0.5 mM isopropyl-β-D-thiogalactoside (IPTG) for SagB and SpdC expression, and 0.2% arabinose for Ulp1 expression. After a 18 h expression, cells were collected by centrifugation and resuspended in buffer containing 50 mM Tris-HCl (pH 7.4), 300 mM NaCl, 1 mM phenylmethylsulfonyl fluoride (PMSF), 1 cOmplete protease inhibitor table (Sigma-Aldrich), 50 μg/ml DNase 1. Resuspended cells were lysed by a 4x passage through an Emulsiflex C3 homogenizer (Avestin) at 15,000 p.s.i. Lysed cells were separated from unbroken cells by centrifugation at 12,000*g*, 4°C for 10 minutes. Membranes were pelleted by ultracentrifugation at 100,000*g*, 4°C for 60 minutes in a Beckman 45Ti rotor. For solubilization, membrane pellets were resuspended in buffer B (50 mM Tris-HCl (7.4), 300 mM NaCl, 10% glycerol), homogenized using an IKA T18 UltraTurrax, and then supplemented with 1% w/v dodecyl-maltoside (DDM; Anatrace). Cell membranes were rocked at 4°C for 1 hour before insoluble cell debris was removed by ultracentrifugation at 100,000xg, 4°C, 30 minutes. Equilibrated Ni-NTA agarose (equilibrated with buffer B supplemented with 10 mM imidazole; Qiagen) was resuspended with solubilized membranes and rocked at 4°C for 1 hour before gravity flow through a column. Following flow-through, resin was washed with 20 column volumes (cv) of buffer B supplemented with 10 mM imidazole and 0.05% DDM and then 20 cv of buffer B with 30 mM imidazole and 0.05% DDM. Protein was eluted with 4 cv buffer B with 200 mM imidazole, 0.05% DDM, and 2 mM CaCl_2_. Elution fractions were then loaded onto a 4 mL M1-anti-Flag antibody affinity resin using gravity flow twice. The resin was then washed with 100 ml buffer containing 50 mM HEPES (pH 7.5), 300 mM NaCl, 10% glycerol, 0.05% DDM, and 2 mM CaCl_2_. Protein was eluted in 20 mM HEPES (pH 7.5), 500 mM NaCl, 20% glycerol, 0.1% DDM supplemented with 5 mM EDTA and 0.2 mg ml^-1^ Flag peptide (Genescript). SagB-SpdC was concentrated using a 50 MWCO concentrator (Amicon) and further purified by size exclusion chromatography on a Sephadex S200 Increase 10/300 GL (GE Healthcare) in buffer (for biochemical reactions, buffer contained 50 mM Tris-HCl (7.4), 300 mM NaCl, 10% glycerol, 0.1% DDM; for crystallography, buffer contained 50 mM Tris-HCl (7.4), 300 mM NaCl, 3% glycerol, 0.02% DDM). For biochemical reconstitutions, SagB-SpdC was concentrated into approximately 2 mg ml^-1^ aliquots, flash-frozen with liquid nitrogen, and stored at −80°C. The attempted purification of SagA-SpdC used a similar protocol, with the expression plasmid containing SagA-His_6_ in place of SagB.

### Expression and purification of *S. aureus* individual glucosaminidases and SpdC

Individual SagB-His_6_, SagA-His_6_, and FLAG-SpdC were expressed and purified in a similar manner as described above with the following modifications. Overnight culture of BL21(DE3) *E. coli* containing the plasmid with SagA, SagB, or SpdC was diluted 1:100 in terrific broth supplemented with 0.1% glucose and 50 μg ml^-1^ carbenicillin. Growth conditions and initial purification steps were similar to as described above, with the exception of using a Ni_NTA resin to bind and purify individual hydrolases and the M1-anti-FLAG resin to bind and purify SpdC. After elution from respective resin, protein was concentrated using a 30 MWCO concentrator and further purified using a Sephadex 200 10/300 GL column using buffer containing 50 mM Tris-HCl (pH 7.4), 300 mM NaCl, 0.1% DDM, 10% glycerol.

For the soluble SagB construct, protein was expressed and purified in the same manner with slight modifications. Overnight cultures of BL21 (DE3) *E. coli* containing a pET_28(b)+ with SagB lacking its transmembrane helix (32-284 aa) was diluted 1:100 in LB. Cells were grown at 37°C at 200 r.p.m. until A_600nm_=0.4 and then cooled to 18°C; at A_600nm_=0.6, protein expression was induced with 0.5 mM IPTG. After 18 hr expression, cells were collected by centrifugation and lysed as described above. After lysis, unbroken cell debris was removed by centrifugation at 12,000*g*, 4°C for 10 minutes. Supernatant was further clarified by ultracentrifugation at 100,000*g*, 4°C for 30 minutes. Ni-NTA resin was equilibrated with clarified supernatant for 1 hour, rocking at 4°C. Following flowthrough, resin was washed with 20 column volumes (cv) of buffer B supplemented with 10 mM imidazole and then 20 cv of buffer B with 30 mM imidazole. Protein was concentrated using a 10 MWCO concentrator tube (Amicon) and then further purified using size exclusion chromatography with Sephadex 75 10/300 GL and a buffer containing 50 mM Tris-HCl (pH 7.4), 300 mM NaCl, 10% glycerol. Aliquots of concentrated soluble SagB at approximately 2-4 mg ml^-1^ were then flash-frozen with liquid nitrogen and stored at −80°C.

### Expression and purification of *S. aureus* SagB with truncated SpdC (1-256 amino acids)

*S. aureus* SagB-SpdC (1-256 amino acids) was expressed and purified in the same manner with slight modifications. Overnight cultures of BL21 (DE3) *E. coli* containing the pDUET with SUMO_Flag_SpdC (1-256 aa) and SagB-His_6_, and an arabinose-inducible Ulp1 protease plasmid (pAM174) were diluted 1:100 in terrific broth supplemented with 0.1% glucose. Cultures were grown at 30°C with shaking at 200 r.p.m. until A_600nm_ = 0.6 and then shifted to 24°C. At an A_600nm_ = 1.1, protein expression was induced by addition of 0.5 mM isopropyl-β-D-thiogalactoside (IPTG) for SagB and SpdC expression, and 0.2% arabinose for Ulp1 expression and grown for 16 h. Purification of SagB-SpdC was similar to as described above.

### Expression and purification of *S. aureus* SagB-SpdC cysteine mutants

The *S. aureus* SagB-SpdC cysteine mutants were expressed and purified in the same manner with slight modifications. Overnight cultures of BL21 (DE3) *E. coli* containing the pDUET with SUMO_Flag_SpdC and SagB-His_6_ with the cysteine mutants, and an arabinose-inducible Ulp1 protease plasmid (pAM174) were diluted 1:100 in terrific broth supplemented with 0.1% glucose. Cultures were grown at 25°C with shaking at 200 r.p.m until A_600nm_ = 0.6 and then shifted to 20°C. At an A_600nm_ = 1.1, protein expression was induced by addition of 0.5 mM isopropyl-β-D-thiogalactoside (IPTG) for SagB and SpdC expression, and 0.2% arabinose for Ulp1 expression and grown for 16 hours. Purification of SagB-SpdC was similar to as described above with the following modifications. Pelleted cells were resuspended in 50 mM Tris-HCl (pH 7.4), 300 mM NaCl, 1 mM phenylmethylsulfonyl fluoride (PMSF), 1 cOmplete protease inhibitor table (Sigma-Aldrich), 50 μg/ml DNase 1. To facilitate disulfide formation, 0.3 mM CuSO4 and 0.3 mM 1,10-phenanthroline were added to the mixture. The remaining purification steps followed those described above.

### PAGE autoradiograph experiments with PG oligomers and *S. aureus* hydrolases

The protocol for analyzing and preparing peptidoglycan oligomers was adapted from similar methods previously reported^17,41^. A PBP2 construct^19^ (59-716 amino acids, 1 μM) was incubated with synthetic-[^14^C]-Lipid II analog^18^ (20 μM in DMSO; specific activity=300 μCi/μmol from UDP-[^14^C]-GlcNAc (American Radiolabeled Chemicals, Inc.) in reaction buffer (50 mM MES (6.5), 100 mM CaCl_2_) with 20% DMSO (v/v). After mixing the Eppendorf tube by flicking and spinning down the reaction mixture, polymerization proceeded at room temperature for 2 hours. To test oligomers prepared from SgtB* (SgtB^Y181D^)^30^, similar conditions were set up with 800 nM SgtB*. Proteins were removed by precipitation by heating the reaction mixture at 95 °C for 10 minutes and then spinning down the precipitated protein. The PG oligomers were then aliquoted into separate Eppendorf tubes (10 μl mixture for each reaction) and a hydrolase (SagA, SagB, SagB-SpdC, or the ΔTM-SagB; 3 μM) was added. For reactions in which the individual hydrolase was incubated with SpdC, the hydrolase SagA or SagB (3 μM) were added to SpdC (3 μM) for 1 hour on ice before addition to the PG mixture. After overnight incubation, the reaction was quenched with heat inactivation (95 °C for 10 minutes) and the reactions were dried completely using a speed vacuum. Reactions were resuspended in 10 μl of SDS loading buffer and loaded onto a 10% acrylamide-Tris gel. The gel was run at 30 mA for 5 h at 4°C; an anode buffer consisted of 100 mM Tris (pH 8.8) and cathode buffer consisted of 100 mM Tris, 100 mM tricine (pH 8.25), 0.1% SDS^41^. Gels were dried on filter paper (19 x 18.5 cm; Biorad) and then exposed to a phosphor screen for at least 24 h. Phosphor screens were imaged using an Azure Sapphire Biomolecular Imager (Azure biosystems). Images were further analyzed using ImageJ.

Reactions to test hydrolase activities with concurrent PBP2 transglycosylase activity were set up and analyzed in a similar manner with some modifications. A reaction mixture was prepared with synthetic-[^14^C]-Lipid II analog^18^ (20 μM in DMSO; specific activity=300 μCi μmol^-1^ from UDP-[^14^C]-GlcNAc (American Radiolabeled Chemicals, Inc.), reaction buffer (50 mM MES (6.5), 100 mM CaCl_2_), 20% DMSO (v/v). PBP2 (59-716 amino acids; 5 μM) and the hydrolase (SagA, SagB, SagB-SpdC; 3 μM) was then added). Similar methods were used to test the activities of interface mutants of SagB-SpdC. To test the activities of disulfide-linked SagB-SpdC complexes, protein under reducing conditions were treated with 5 mM DTT on ice for 30 minutes before the addition to the reaction mixture that included the addition of 5 mM DTT. After overnight incubation, the reaction was quenched with heat inactivation (95 °C for 10 minutes) and the reactions were dried completely using a speed vacuum. Reactions were resuspended in 10 μl of 2x SDS loading buffer and loaded onto a 10% acrylamide-Tris gel. The gel was run and analyzed as described above.

Testing hydrolase activities with wall-teichoic acid labeled peptidoglycan oligomers was adapted from similar methods previously reported^36^. Peptidoglycan oligomers were prepared as described above although with unlabeled, synthetic-Lipid II (20 μM). After precipitation to remove the PGT, a portion of the mixture was incubated with TagT (1 μM) and [^14^C]-LII_A_^WTA^ (8 μM) for four hours at room temperature. The TagT ligase was then heat inactivated, precipitated, and removed from the reaction mixture. The resulting wall teichoic acid-labeled oligomers were then incubated with SagB (3 μM), SagB-SpdC (3 μM), or mutanolysin (2.5 U ml^-1^). The remaining portion of unlabeled PG oligomers was also incubated with the SagB-SpdC complex (3 μM). After an overnight incubation at room temperature, the reaction was quenched with heat inactivation (95 °C for 10 minutes). To test TagT ligation after SagB-SpdC incubation, TagT (1 μM) and [^14^C]-LII_A_^WTA^ (8 μM) was added to the respective reaction for four additional hours. Reactions were dried completely using a speed vacuum and the mixture was re-dissolved in 10 μl of 2x SDS loading buffer and loaded onto a 10% acrylamide-Tris gel. The gel was run and analyzed as described above.

### Western blot analysis of hydrolase activities with peptidoglycan oligomers prepared from *S. aureus* Lipid II

The protocol for detecting peptidoglycan oligomers prepared from extracted *S. aureus* Lipid II was adopted from similar methods previously reported^19,23,29^. To generate uncrosslinked PG oligomers, *S. aureus* Lipid II (10 μM) was added to reaction buffer (50 mM MES (6.5), 10 mM CaCl_2_), 20% DMSO (v/v) and thetranspeptidase inactive construct PBP2^S398G^ (5 μM) was added to the reaction. To test ongoing hydrolase and polymerase activity, the reaction mixture was aliquoted into separate tubes and the respective glucosaminidase was also added (SagA, SagB, or SagB-SpdC; 3 μM). For a mutanolysin-digest reaction, 2.5 U ml^-1^ of mutanolysin (Sigma-Aldrich) was added^42^. If testing hydrolase activity with pre-assembled PG oligomers, PBP2^S398G^ was precipitated after heat inactivation at 95°C. PG oligomers were aliquoted into separate 10 μl reactions, and the hydrolase was added as described above. Reactions were inactivated at 95°C for 10 minutes. To label uncrosslinkedPG oligomers, BDL (4 mM in H_2_O) and *E. faecalis* PBPX^31^ (20 μM) was added to the reaction mixtures. After a 1 h incubation at room temperature, 2x loading buffer was added to quench the reactions. To generate crosslinked PG, *S. aureus* Lipid II (10 μM) was added to reaction buffer (50 mM MES (6.5), 10 mM CaCl_2_), 20% DMSO (v/v), BDL (4 mM), and a wild type PBP2 construct (5 μM). After the addition of PBP2, the respective glucosaminidase was also added (SagB or SagB-SpdC; 3 μM) and the reactions were incubated at room temperature for 5 hours. Reactions were split into two aliquots after heat inactivation at 95°C, and lysostaphin (100 μg ml^-1^) was added to one aliquot and then incubated at 37°C for 2 hours. 2x loading dye was added to each reaction, and these mixtures were loadedonto a 4-20% polyacrylamide gel which ran at 175 V for 1 h. The gel was transferred to PVDF membrane (Biorad) at 10 mV for 1 h. After incubating with starting block (Thermo Scientific, catalog number #37578), the membrane was rocked with HRP-streptavidin (1:5000) in TBS-T and then washed repetitively. The membrane was imaged using the chemiluminescence function on an Azure imager (Azure biosystems). Images were further analyzed using ImageJ.

### LC-MS method for detecting digest products of *S. aureus* glucosaminidases and mutanolysin

The cleaved muropeptide products of glucosaminidase reactions (SagA, SagB, SagB-SpdC) were analyzed by adopting a previously reported LC-MS method for detecting mutanolysin-digested products^42^. Hydrolase reactions were prepared using synthetic Lipid II (20 μM), reaction buffer (50 mM MES (6.5), 10 mM CaCl_2_), 20% DMSO (v/v), PBP2S398 (5 μM), and then the respective hydrolase (SagA, SagB, SagB-SpdC, 3 μM) in a total volume of 50 μl. A control mutanolysin reaction was likewise set-up with 5 U ml^-1^ mutanolysin. After an overnight incubation, sodium borohydride (10 mg ml^-1^ in H_2_O; equal reaction volume) was added and incubated for 20 minutes at room temperature. The reaction was quenched with the addition of 20% phosphoric acid which also adjusted the pH to 3-4. Reactions were completely dried under a nitrogen stream and then resuspended in H_2_O. LC-MS analysis was conducted using an Agilent Technologies 1200 series HPLC in line with an Agilent 6210 TOF mass spectrometer with electrospray ionization (ESI) and operating in positive mode. Muropeptide cleavage products were separated using a Waters Symmetry Shield RP18 column (5 μm, 3.9 x 150 mM) with a matching guard column and the following method: 0.4 mL min^-1^ solvent A (water/0.1% formic acid) for 5 minutes followed by a linear gradient of 0 to 40% solvent B (acetonitrile/0.1% formic acid) over 25 min. Mass spectrometry data was analyzed using Agilent MassHunter Workstation Qualitative Analysis software version B.06.00 and Prism 7.0b.

### LC-MS method for detecting lipid-linked PG oligomers

To character lipid-linked cleavage products, we adapted previously published methods^23^. Peptidoglycan oligomers were prepared by incubating Lipid II (40 μM) with PBP2^S398G^ (5 μM) in 20% DMSO and reaction buffer (50 mM MES (6.5), 100 mM CaCl_2_) and 20% DMSO (v/v) for a total of 2 h. SagB-SpdC (3 μM) was added to the mixture and then incubated for approximately 8 hours. Enzymes were heat inactivated at 95°C for 5 minutes. After cooling reactions to room temperature, ColM (1 mg mL^-1^) was added to the mixture and incubated for approximately 3 hours at room temperature^23^. Protein was precipitated with equal volume methanol. Dried reactions were then resuspended in 20 μL of H_2_O.

### Glycan strand assay

Overnight cultures of wild-type HG003 *S. aureus,* SHM056, and JP132 were diluted 1:100 into 1 L of TSB each and grown at 30 °C for 5 hours. Cells were harvested and sacculi were isolated following a previously published protocol^43^ with the modification that samples were boiled in SDS for 3 hours. Sacculi were resuspended to 5 mg/mL in 500 μL of 25 mM NaH_2_PO_4_ pH 7.0 and treated with 100 μg/mL lysostaphin at 37 °C with shaking for 9 hours. LytA was then added to 100 μg/mL and shaking continued at 37 °C for another 14 hours. Enzymes were heat inactivated at 95 °C for 5 min. Proteins were precipitated with an equal volume of MeOH and removed by centrifugation. To label glycans, we adapted methods previously published^44^. Dried samples were resuspended in 25 μL of 1 M 2-methylpyridine borane complex in DMSO and 25 μL of 200 mM 8-aminonaphthalene-1,3,6-trisulfonic acid in 15% acetic acid. Reactions were incubated at room temperature overnight protected from light. Reactions were quenched with 450 μL of H_2_O at room temperature for 1 hr, and an equal volume of MeOH was added. After spinning down, the supernatants were removed to new tubes and dried. Dried samples were resuspended in 50 μL of 1x loading buffer (125 mM Tris-tricine pH 8.2, 10% glycerol) and loaded on a 20% polyacrylamide gel, which ran for 9 hours at 25 mA. The gel was visualized under UV light at 365 nm.

### Crystallization of SagB-SpdC^1-256^ and data collection

For crystallization trials, SagB-SpdC^1-256^ was purified as described above. Freshly purified SagB-SpdC^1-256^ was concentrated to 35-40 mg/ml and immediately reconstituted into lipidic cubic phase by mixing protein and monoolein (Hampton Research) at a 1:1.5 ratio by mass, using the coupled syringe method^27^. All samples were mixed at least 100 times prior to crystallization trials. The resulting mixture was dispensed onto glass plates in 35-50 nL drops, then overlaid with 600 nL precipitant solution using an NT8 robot (Formulatrix). Crystals of SagB-SpdC^1-256^ were grown in precipitant solution containing 24-32% PEG400, 500 mM (NH_4_)_2_SO_4_, and 100mM sodium acetate or sodium citrate pH 4.4-5.0; at higher pH crystallization required higher concentrations of PEG400. Most crystals appeared within 36 hours of drop setting and were full-grown in 3-7 days. Crystals were harvested using mesh loops and flash-frozen in liquid nitrogen.

Diffraction data were collected at Argonne National Laboratory using NE-CAT beamlines 24-ID-C and 24-ID-E. Two rounds of grid scanning with large and then small beam size were used to locate crystals on the mesh and then precisely determine their positions. All data were collected at 0.979 Å. Datasets were collected 0.2-s exposure and a 0.2° oscillation angle. The presence of multiple crystals on the mesh prevented the collection of full datasets from individual crystals. Data were indexed and integrated in XDS^45^; the SagB-SpdC crystals belonged to the C2 space group. Diffraction data was processed using structural biology software accessed through the SBGrid consortium^46^. Partial datasets from five well-diffracting crystals were then scaled and merged using the CCP4 suite^47^ program AIMLESS^48^.

### Structure determination and refinement

The structure was determined by molecular replacement in Phaser using an unpublished structure of the soluble domain of SagB (PDB# 6FXP)^20,49^. We attempted to place SpdC by molecular replacement using models prepared from the structure of *Methanococcus maripaludis* CAAX protease Rce1 (PDB# 4CAD, 20% sequence identity to SpdC)^50^, but no good solutions were found. Following placement of soluble SagB, only weak density was visible for the transmembrane helices of the complex; those that were clearest were initially modeled as poly-alanine helices using Coot^51,52^. After one round of automated refinement using phenix.refine^53^, placement of all nine transmembrane helices was possible, but residue identities were not obvious. Initial assignment was made by sequence alignment and comparison to the structure of Rce1, which appeared to have a similarly threaded transmembrane domain. Density for sidechains in the transmembrane domain became much clearer after several rounds of manual building and automated refinement, and simulated annealing composite omit maps were used to check for model bias and correct errors in the register throughout the process. Towards the end of refinement, a region of clear density remained present at a crystallographic interface between two copies of SagB near residues 38-45 and appeared to be a peptide forming a continuous β-sheet with the two copies of SagB. This peptide is on the unit cell edge but does not appear to be part of either protein; we think it is likely FLAG peptide that carried through the purification and may orient either direction at this interface. Water molecules, sulfate ions, PEG molecules were also added near the end of refinement. Structure quality was assessed using MolProbity^54^, and figures were prepared using PyMOL.

## Notes

### Competing Interest Statement

The authors have declared no competing interest.

