## Supplementary Information for "Structure and reconstitution of a hydrolase complex that releases peptidoglycan from the membrane after polymerization"

| Beta lactam treatment | Gene number | Gene name | Corrected p-value | Ratio |
|---|---|---|---|---|
| Oxacillin | *USA300HOU_RS09425* | *sagB* | 1.48E-13 | 0.0358 |
| | *USA300HOU_RS12635* | *spdC* | 5.26E-17 | 0.0599 |
| Cefoxitin | *USA300HOU_RS09425* | *sagB* | 1.84E-06 | 5.0994 |
| | *USA300HOU_RS12635* | *spdC* | 4.48E-11 | 6.2723 |
| Mecillinam | *USA300HOU_RS09425* | *sagB* | 6.16E-07 | 0.1675 |
| | *USA300HOU_RS12635* | *spdC* | 4.19E-06 | 0.2869 |

**Supplementary Table 1. Treatment of a methicillin-resistant *Staphylococcus aureus* transposon library with sublethal concentrations of beta-lactams identified *sagB* and *spdC* as the only two genes with a unique cell wall phenotype.**

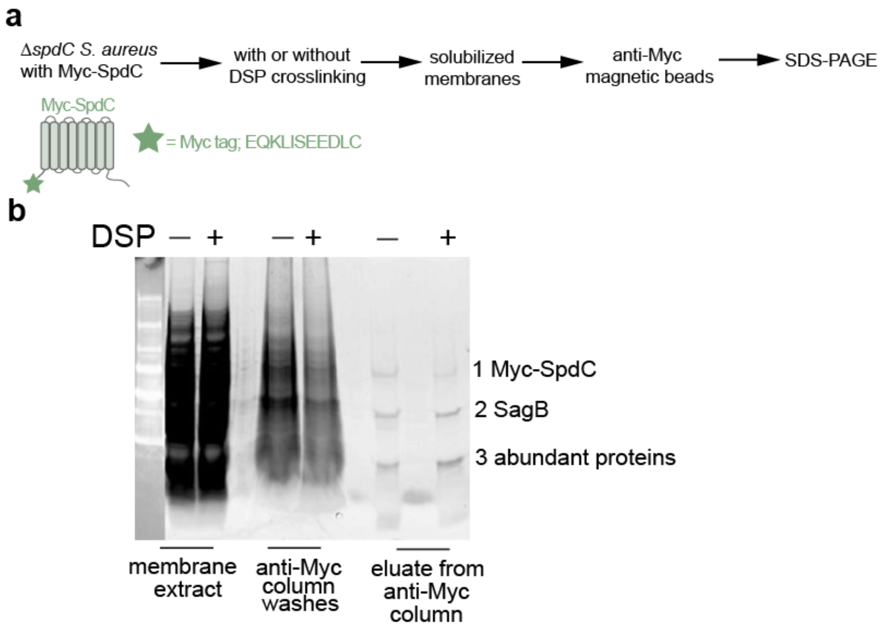


**Supplementary Figure 1. The interaction between SagB and SpdC was first observed by co-immunoprecipitation of SagB with a Myc-epitope tagged SpdC in a *S. aureus* Δ*spdC* background.** Untreated cells and cells treated with a chemical crosslinker, DSP (dithiobis(succinimidyl propionate); Thermo Scientific, Catalog # 22585), were lysed and their membranes isolated. Solubilized cell membranes were applied to anti-myc magnetic beads, which were washed extensively before bound proteins were eluted and then analyzed using SDS-PAGE. Protein bands from the sample without crosslinker (1-3) were then subjected to LC-MS-MS analysis, which identified the middle band as SagB (Supplementary Table 2).

Band 1:

| Unique peptides | Total peptides | Reference | Gene symbol | MW (kDa) |
|---|---|---|---|---|
| 35 | 102 | P02976_SPA_STAA8 | *spa* | 56.40 |
| 16 | 17 | Q2FY15_Q2FY15_STAA8 | *SAOUHSC_01659* | 51.05 |
| 14 | 15 | Q2FWW1_Q2FWW1_STAA8 | *SAOUHSC_02161* | 65.53 |
| 13 | 14 | Q2FWF0_ATPB_STAA8 | *atpD* | 51.37 |
| 13 | 14 | Q2FWH5_Y2316_STAA8 | *SAOUHSC_02316* | 56.91 |
| 13 | 13 | Q2FYZ4_GLPD_STAA8 | *glpD* | 62.35 |
| 12 | 14 | Q2FWE8_ATPA_STAA8 | *atpA* | 54.55 |
| 12 | 12 | Q2FV16_Q2FV16_STAA8 | *mqo* | 55.96 |
| 11 | 21 | Q2FVT1_LYRA_STAA8 | ***spdC*** | 46.76 |
| 10 | 10 | Q2G0N0_EFTU_STAA8 | *tuf* | 43.08 |

Band 2:

| Unique peptides | Total peptides | Reference | Gene symbol | MW (kDa) |
|---|---|---|---|---|
| 15 | 30 | P60430_RL2_STAA8 | *rplB* | 30.14 |
| 12 | 16 | Q2FWW9_Q2FWW9_STAA8 | *SAOUHSC_02152* | 32.93 |
| 12 | 13 | Q2FXF4_Q2FXF4_STAA8 | ***sagB*** | 32.49 |
| 11 | 11 | Q2G2S6_PRSA_STAA8 | *prsA* | 35.62 |
| 10 | 10 | Q2FXZ9_Y1676_STAA8 | *SAOUHSC_01676* | 35.16 |
| 10 | 10 | Q2G2D8_Q2G2D8_STAA8 | *SAOUHSC_00634* | 35.05 |
| 9 | 15 | Q2FW06_RL3_STAA8 | *rplC* | 23.7 |
| 8 | 14 | Q2G0B5_Q2G0B5_STAA8 | *SAOUHSC_00690* | 26.67 |
| 8 | 8 | Q2FVL2_Q2FVL2_STAA8 | *SAOUHSC_02699* | 28.89 |
| 8 | 8 | Q2G0N0_EFTU_STAA8 | *tuf* | 43.08 |

Band 3:

| Unique peptides | Total peptides | Reference | Gene symbol | MW (kDa) |
|---|---|---|---|---|
| 10 | 10 | Q2G0F2_Q2G0F2_STAA8 | *SAOUHSC_00617* | 18.58 |
| 9 | 29 | Q2FXS8_RL21_STAA8 | *rplU* | 11.33 |
| 9 | 9 | Q2FW30_RS13_STAA8 | *rpsM* | 13.71 |
| 8 | 16 | Q2FXQ1_RL20_STAA8 | *rplT* | 13.68 |
| 7 | 8 | P0A0F8_RL15_STAA8 | *rplO* | 15.59 |
| 7 | 8 | Q2G247_Y1855_STAA8 | *SAOUHSC_01855* | 17.99 |
| 7 | 7 | Q2FW11_RL22_STAA8 | *rplV* | 12.83 |
| 7 | 7 | Q2FVN6_Q2FVN6_STAA8 | *SAOUHSC_02666* | 13.33 |
| 6 | 7 | Q2FW23_RS5_STAA8 | *rpsE* | 17.73 |
| 6 | 6 | Q2FXM1_Q2FXM1_STAA8 | *SAOUHSC_01814* | 15.22 |

**Supplementary Table 2. List of proteins identified by LC-MS-MS analysis of the sample obtained from immunoprecipitation of Myc-SpdC.** Each protein band shown in Supplementary Fig. 1 was analyzed separately. For each of the three bands observed in SDS-PAGE, here is the list of the top ten proteins with the highest coverage in LC-MS-MS analysis.

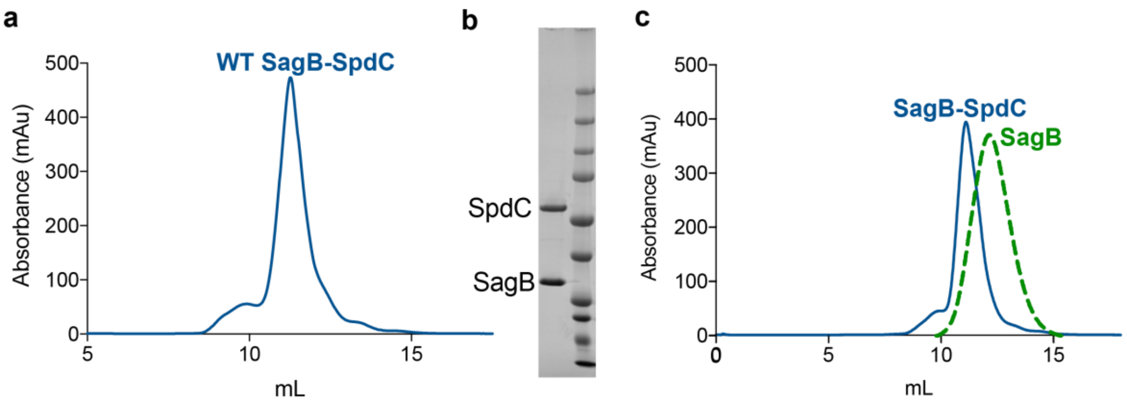


**Supplementary Figure 2. Co-purification of SagB and SpdC results in a stable SagB-SpdC complex. a,** An expression plasmid containing an N-terminal SUMO-FLAG epitope tag on SpdC and a C-terminal hexa-histidine tag on SagB was transformed into *E. coli* BL21-DE3 that also contained an arabinose-inducible Ulp1 protease plasmid. Purification steps included two affinity columns: nickel resin to pull down SagB-$His_6$ and an anti-FLAG M1 antibody resin to pull down an amino-FLAG-SpdC (SUMO removed by Ulp1 during co-expression). Eluate fractions were then concentrated and further purified by size exclusion chromatography with an Increase Sephadex S200 column 10/300 GL. **b,** SDS-PAGE of the dominant peak, which corresponds to the correct molecular weight of the DDM micelle, SagB, and SpdC, shows the presence of SpdC and SagB. **c,** Overlaid size exclusion chromatograms from a SagB-SpdC co-purification and a full-length SagB purification show a peak shift to a higher molecular weight in the presence of SpdC.

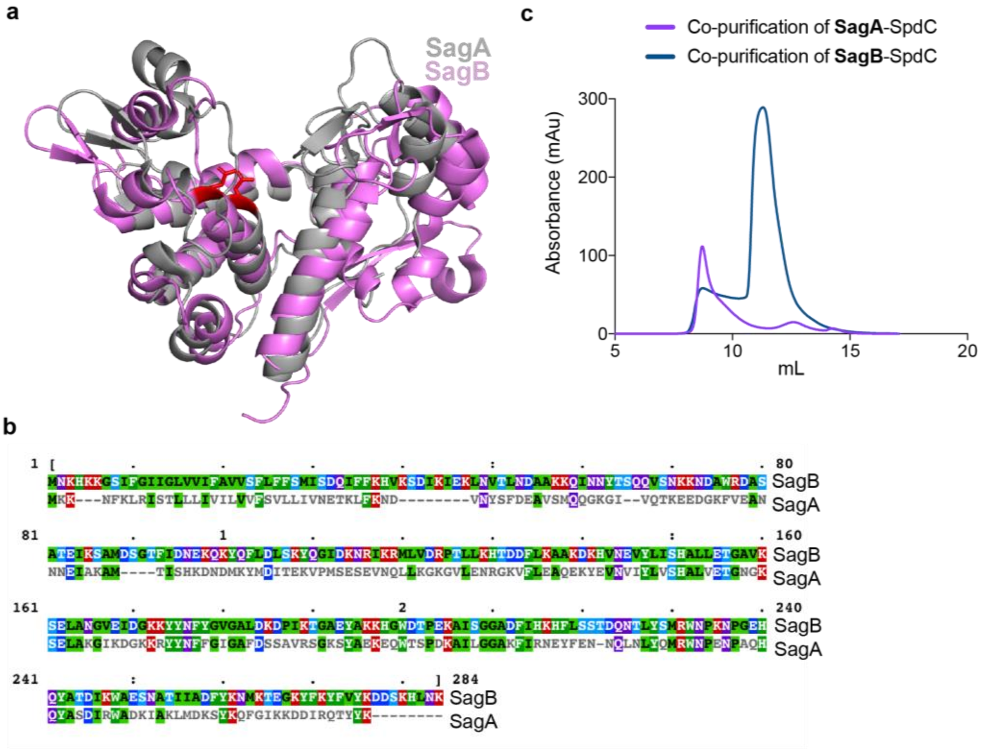


**Supplementary Figure 3. SagB is related to the other *S. aureus* membrane-anchored glucosaminidase, SagA, but SagA does not form a stable complex with SpdC. a,** Alignment of SagA and SagB glucosaminidase domains, colored gray and magenta, respectively (PDB# 4PIA and 6FXP)[4,5]. **b,** Primary sequence alignment of SagA (bottom) and SagB, with residues colored by type. Alignments were made using ClustalO and then coloured in MView (https://www.ebi.ac.uk/Tools/msa/clustalo/). SagA and SagB are 53% similar. **c,** Parallel attempts were made to co-purify SagB-SpdC and SagA-SpdC using FLAG-SpdC and histidine-tagged SagA or SagB. Solubilized membranes were purified on nickel resin and then anti-FLAG resin. Eluate from anti-FLAG resin was concentrated and run through a size exclusion column. A stable SagB-SpdC complex was obtained (blue trace), whereas SagA (violet trace) did not co-elute from anti-FLAG resin with FLAG-SpdC.

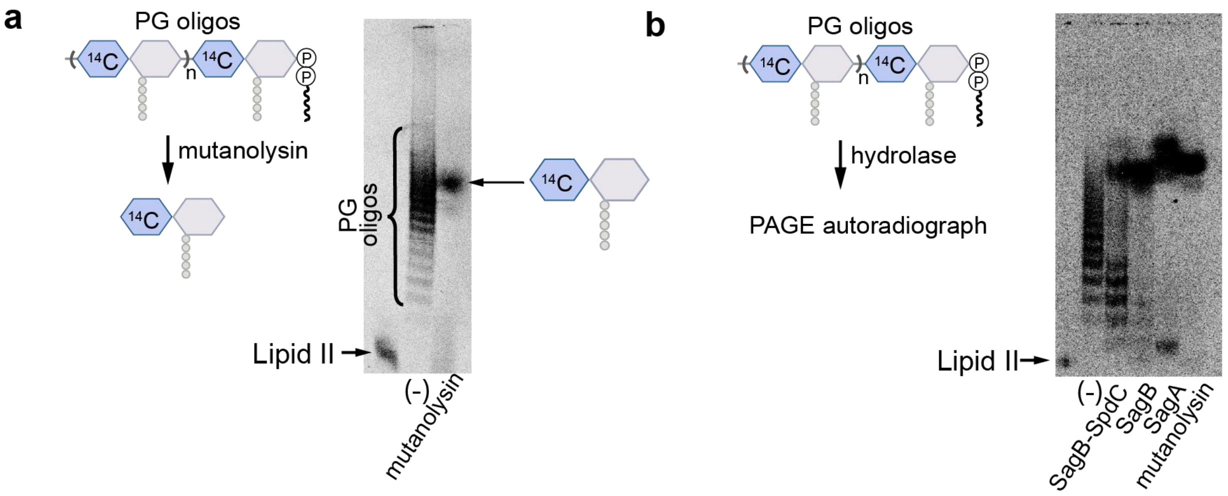


**Supplementary Figure 4. Muramidase cleavage of radiolabeled peptidoglycan oligomers results in a diffuse signal at the same approximate position as cleavage products of glucosaminidases. a,** Mutanolysin is a well-characterized muramidase that cleaves peptidoglycan strands into disaccharide units[1]. Radiolabeled peptidoglycan oligomers run as a ladder (lane 2), but incubation with mutanolysin (lane 3) changes the ladder to a diffuse signal that migrates towards the middle of the gel. **b,** Radiolabeled Lipid II incubated with SgtB* (SgtB$^{Y181D}$) resulted in a ladder of PG oligomers as visualized in the PAGE autoradiograph[2,3]. Radiolabeled product bands are similar to hydrolase reactions with PG oligomers prepared using PBP2 (see main text, Fig. 2b). PG oligomer treatment with SagB-SpdC resulted in two discrete signals on the autoradiograph, a ladder at the bottom of the gel and a signal towards the top of the gel that was also present in SagB, SagA, and mutanolysin reactions.

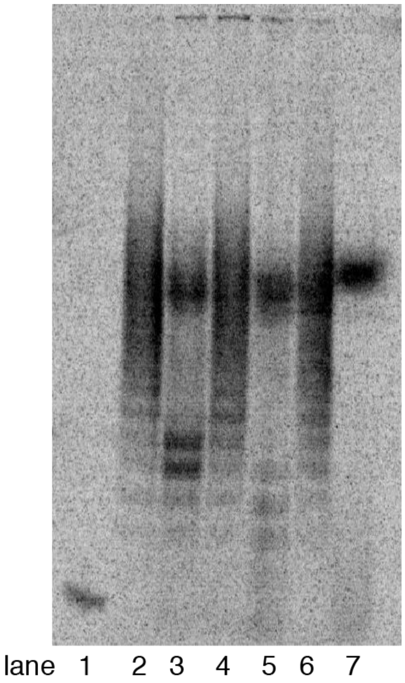


lane 1: [$^{14}$C]-Lipid II
lane 2: [$^{14}$C]-PG oligomers
lane 3: [$^{14}$C]-PG oligomers + SagB-SpdC
lane 4: [$^{14}$C]-PG oligomers + SagB(E155A)-SpdC
lane 5: [$^{14}$C]-PG oligomers + full length SagB
lane 6: [$^{14}$C]-PG oligomers + ΔTM-SagB
lane 7: [$^{14}$C]-PG oligomers + SagA

**Supplementary Figure 5. Hydrolase activities are unaffected by acidic conditions.** Peptidoglycan oligomers were prepared at pH 5.5 in sodium citrate buffer, and then incubated with the respective hydrolase. Cleavage patterns are similar to those observed at pH 6.5 (see main text Fig. 2).

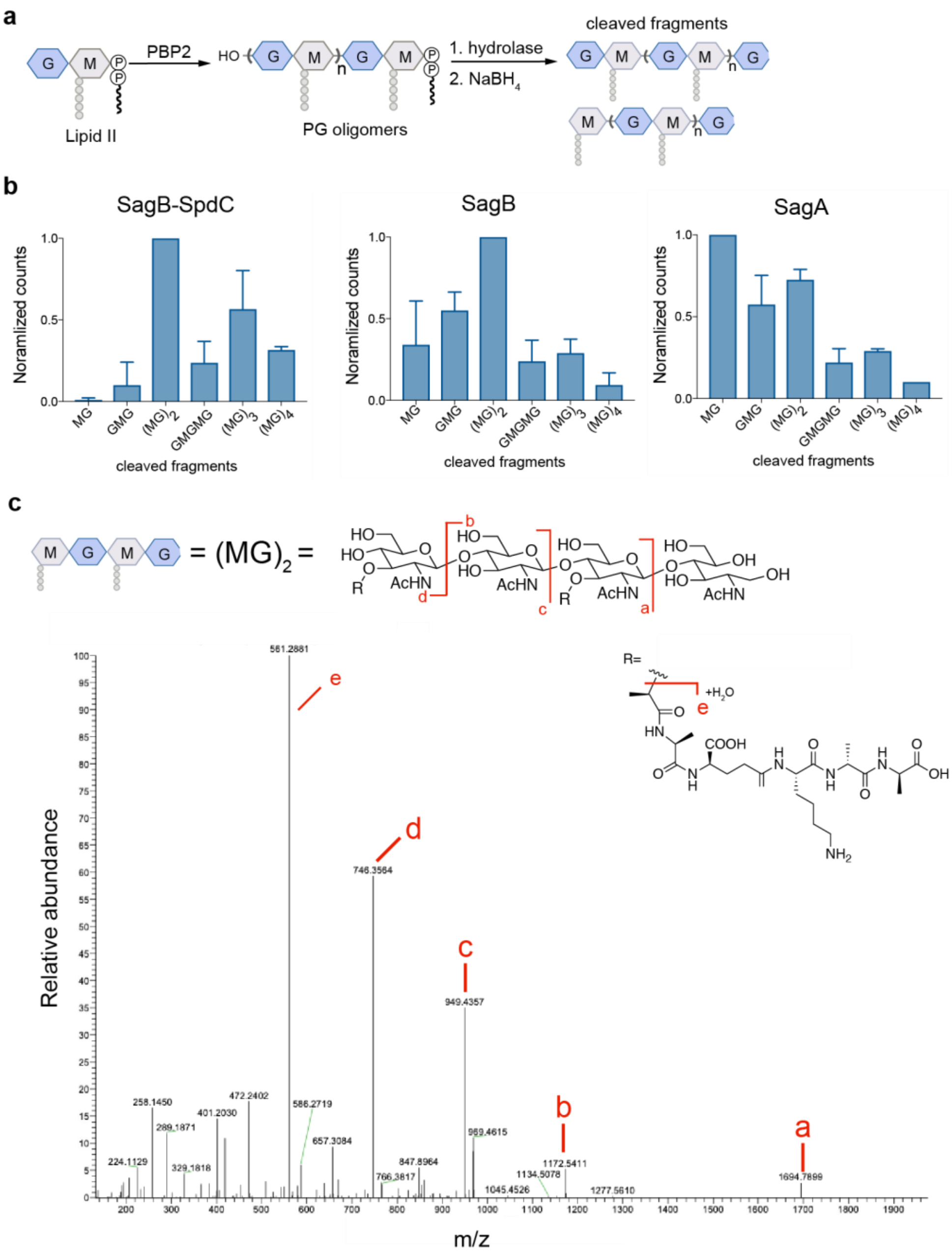


**Supplementary Figure 6. SagB, SagA, and the SagB-SpdC complex cleave peptidoglycan strands into oligosaccharide fragments, but the complex favors longer products**. **a**, This scheme shows the isolation of cleaved peptidoglycan fragments for LC-MS analysis. Peptidoglycan oligomers were incubated with a glucosaminidase (SagA, SagB, or SagB-SpdC). After the cleavage reaction, the reaction mixture was treated with sodium borohydride to reduce any lipid-free fragments[1]. **b**, Molecular ions corresponding to the expected muropeptides were extracted. Integrated peak areas for the muropeptides were normalized relative to one. The bars represent the averages of three biological experiments and the error bar represents standard deviation. These products presumably correspond to the diffuse signal and slower migrating products visualized in the PAGE autoradiograph experiments (see main text Fig. 2). These structures are consistent with glucosaminidase cleavage and also confirm the endolytic cleavage of nascent peptidoglycan oligomers. **c,** Targeted MS-MS of the tetrasaccharide cleavage product $(MG)_2$ from the SagB-SpdC reaction confirms that a GlcNAc is present on the reducing end. The $[M]^{+1}$ was targeted for fragmentation.

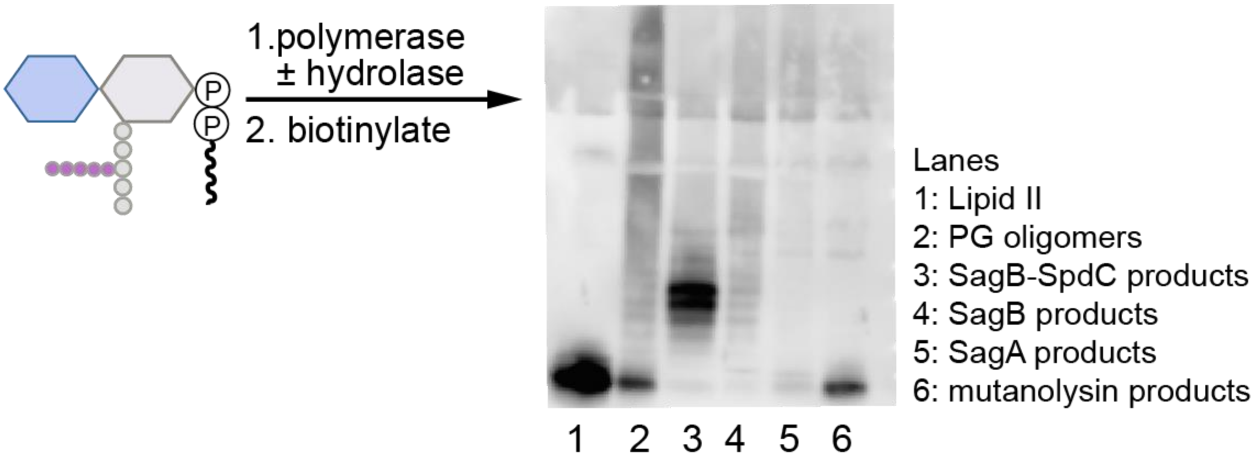


**Supplementary Figure 7. The complex cleaves native PG oligomers to defined lengths similarly to synthetic PG oligomers.** To prepare linear glycan strands, extracted *S. aureus* Lipid II was incubated with transpeptidase-inactive PBP2$^{S398G}$ and then the SagB-SpdC complex or individual hydrolases (SagB or SagA). Reactions were quenched and products were then labeled with biotin-D-lysine (BDL) in the presence of the *E. faecalis* PBPX and visualized using a western blot method with HRP-streptavidin detection[6-8].

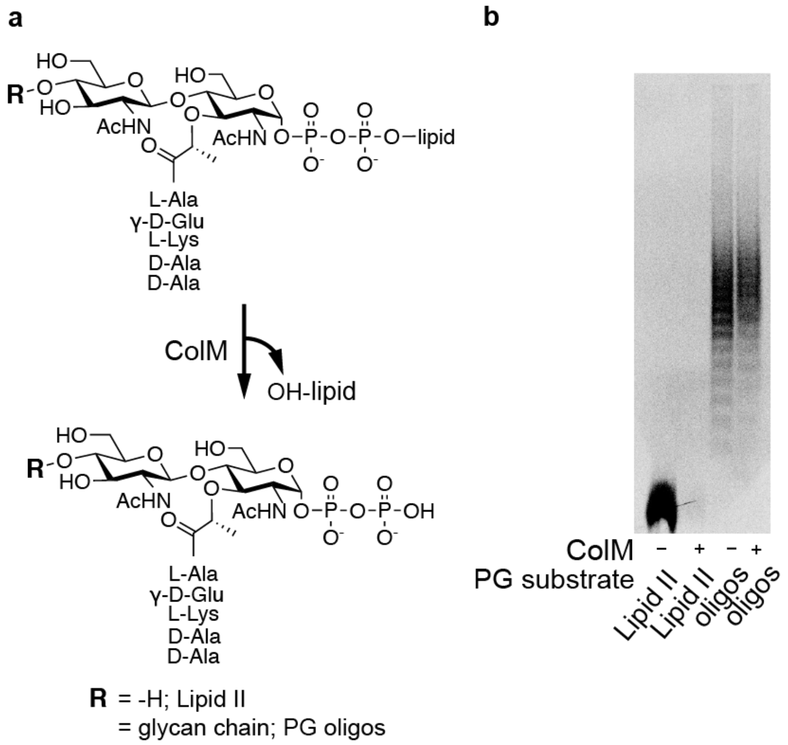


**Supplementary Figure 8. Colicin M (ColM) cleaves the lipid anchor from Lipid II and PG oligomers.** Radiobeled Lipid II runs as a single band at the bottom of PAGE autoradiograph; the signal disappears upon treatment with ColM presumably due to its inability to stay within the gel matrix. Radiolabeled PG oligomers run as a discrete ladder. When incubated with ColM, the oligomers also react but the products migrate similarly to the longer oligos (compare lane 3 to lane 4). Mass spectrometry confirmed the reactions (Supplementary Fig. 9).

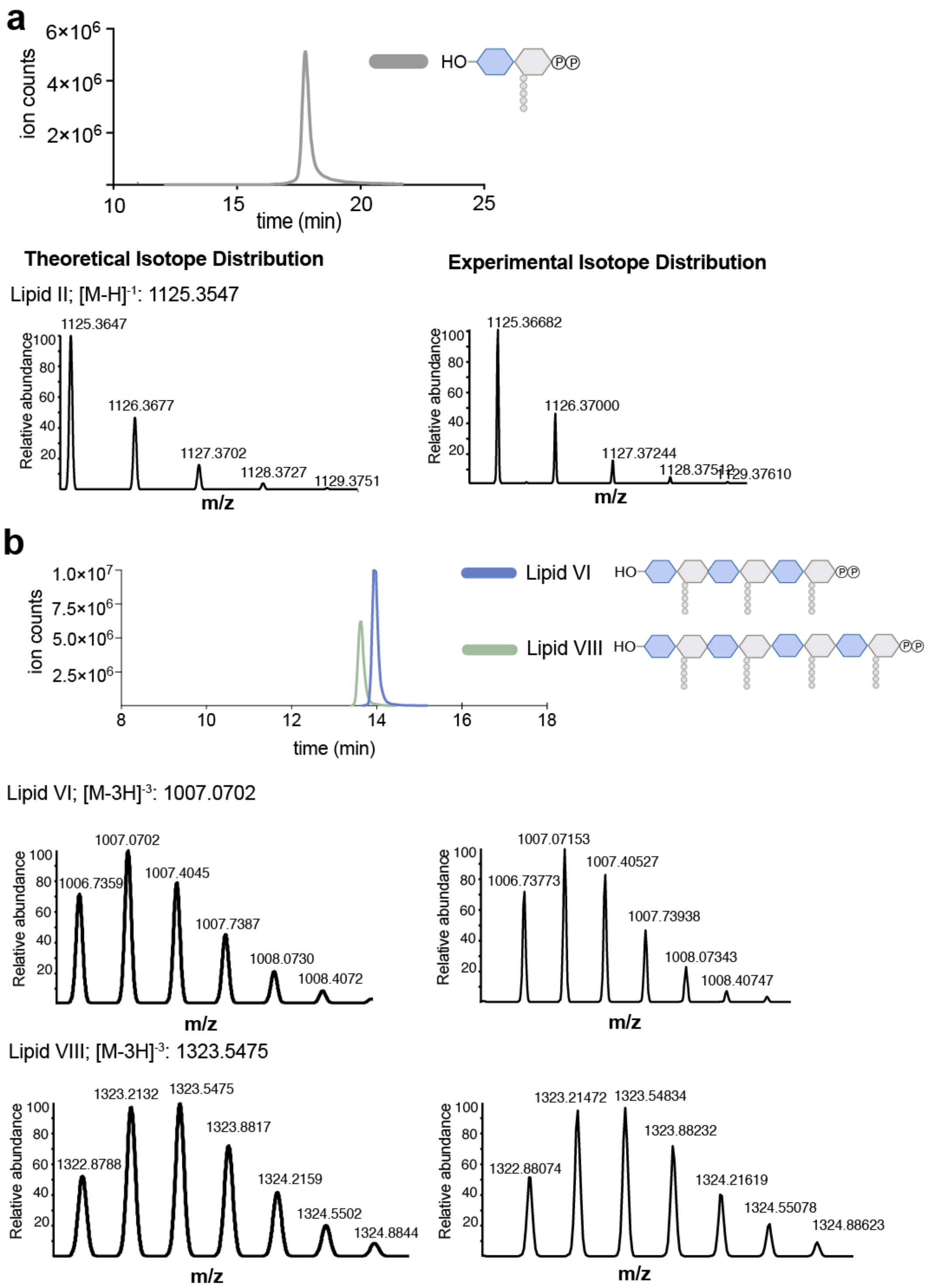


**Supplementary Figure 9: ColM cleaves both Lipid II and linear peptidoglycan strands.**

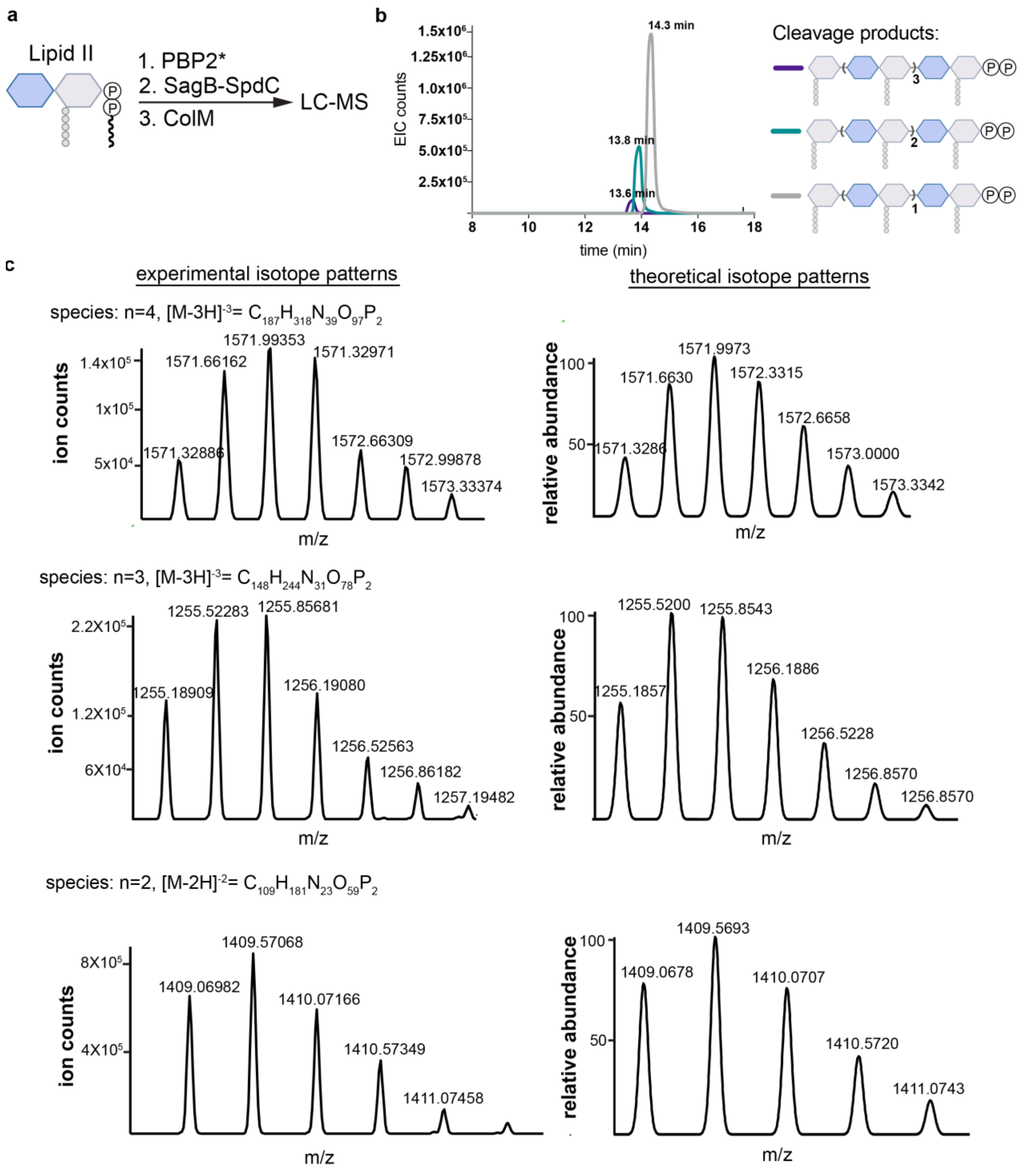


**Supplementary Figure 10. LC-MS analysis of SagB-SpdC reactions incubated with colicin M (colM) confirms the identity of short, lipid-linked cleavage products.** Pre-assembled peptidoglycan oligomers were incubated with SagB-SpdC, and then treated with ColM. Chromatogram traces and isotope patterns for three cleaved peptidoglycan oligomers containing a diphosphate indicate that ColM cleaved the lipid anchor. The odd number of sugars is consistent with glucosaminidase activity.

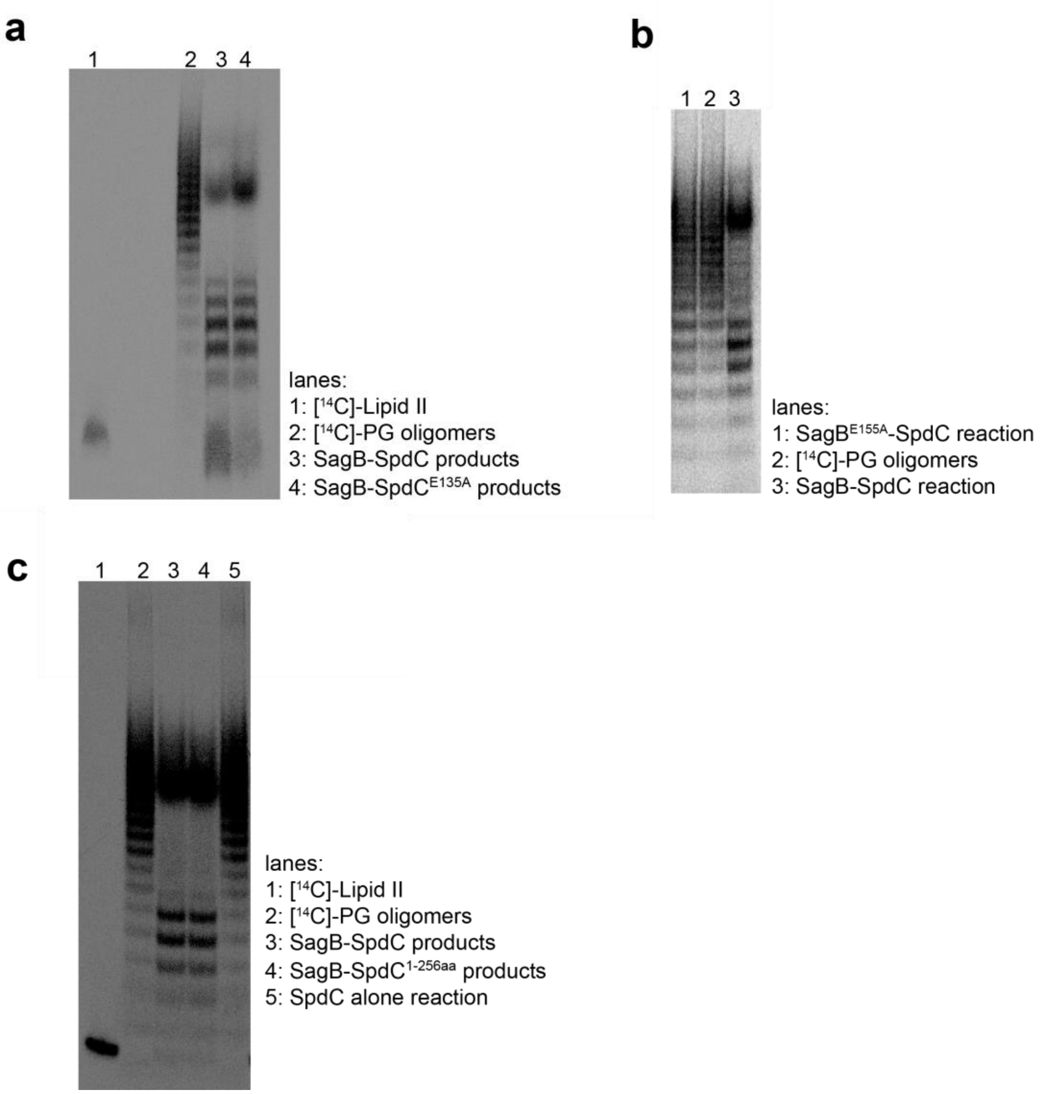


**Supplementary Figure 11. The cleavage products of SagB-SpdC require the catalytic glutamate in SagB, but not the cytoplasmic C-terminal part or the CAAX proteolytic glutamate of SpdC. a,** Activity of wild-type SagB-SpdC in the presence of radiolabeled peptidoglycan oligomers was tested alongside SagB-SpdC$^{E135A}$, in which the proposed CAAX proteolytic active site glutamate[9] is mutated to an alanine. Similar cleavage products are observed, indicating that the conserved SpdC residue is not important for SagB-SpdC glucosaminidase activity. **b,** In the presence of radiolabeled peptidoglycan oligomers, the activity of wild-type SagB-SpdC was compared to the activity of a variant predicted to be catalytically-inactive[10], SagB$^{E155A}$-SpdC. The similarity between the SagB$^{E155A}$-SpdC reaction (lane 1) and control peptidoglycan oligomers (lane 2) indicates that E155 is critical for cleavage. **c,** Radiolabeled peptidoglycan oligomers (lane 2) were incubated with wild-type SagB-SpdC (lane 3), truncated SagB-SpdC$^{1-256}$, which lacks the C-terminal, cytoplasmic region of SpdC (lane 4), and SpdC alone (lane 5). Purification of SagB-SpdC$^{1-256}$ is shown in Supplementary Fig. 16. SpdC alone does not affect the peptidoglycan oligomers, and the cytoplasmic region of SpdC is not required for the activity of the complex.

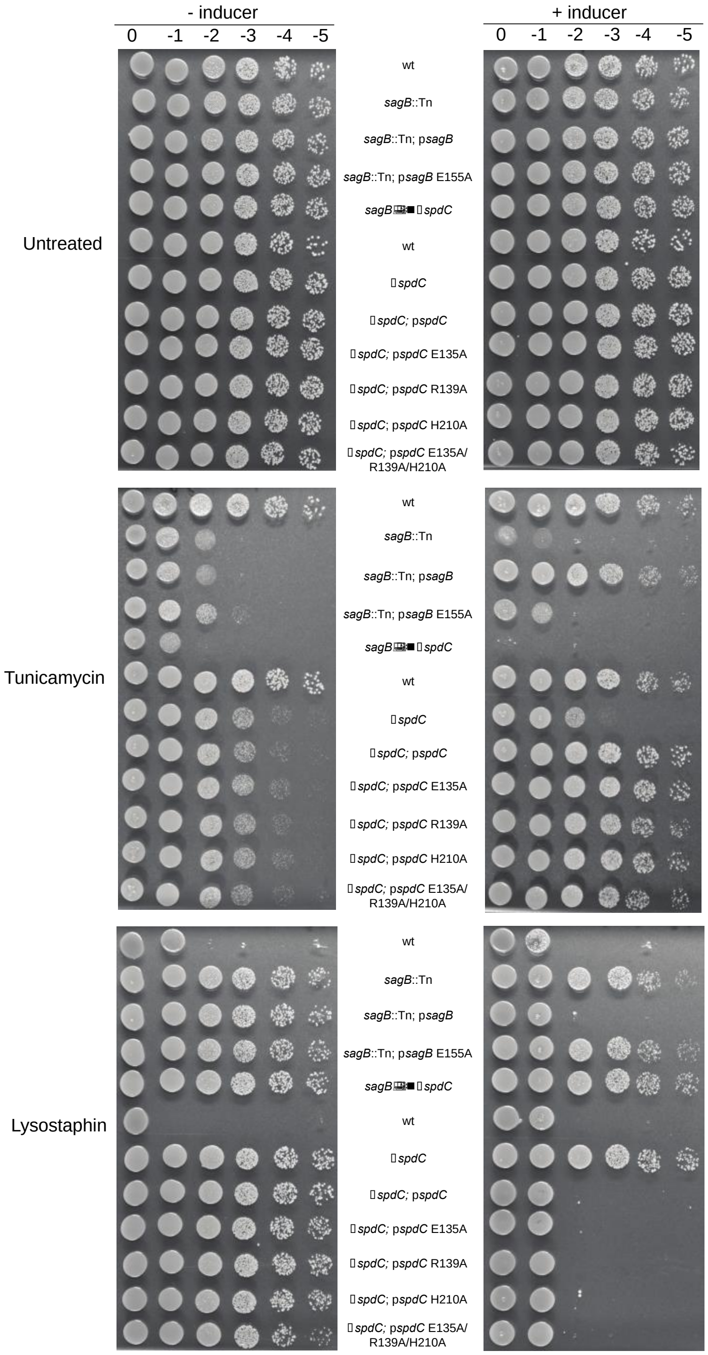


**Supplementary Figure 12. The catalytic glutamate of SagB (E155), but not conserved CAAX protease residues in SpdC (E135, R139, H210)[5], is necessary to complement shared cell wall phenotypes of *sagB* and *spdC* mutants.** Spot dilution series of *S. aureus* strains with mutations in *sagB* or *spdC* were plated on TSA or TSA containing tunicamycin or lysostaphin. Plates shown on the right are replicates of those on the left but the agar contains the inducer, anhydrotetracycline.

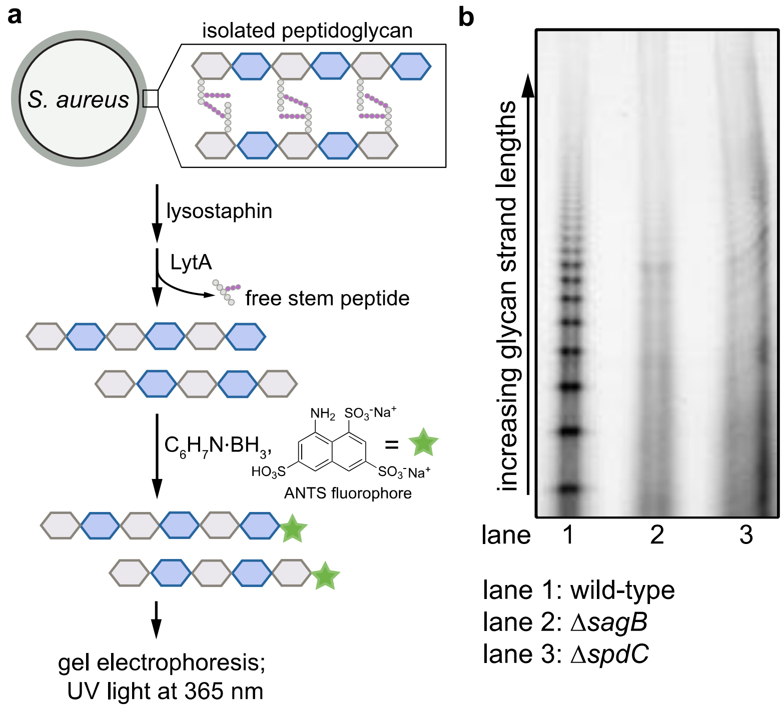


**Supplementary Figure 13. Glycan strand lengths are longer in the absence of *spdC* or *sagB*. a**, Sacculi were isolated from wild-type, Δ*sagB*, and Δ*spdC S. aureus* strains. The purified peptidoglycan was then treated with the endopeptidase lysostaphin, which cleaves the PG crosslinks, and the amidase LytA, which removes the remaining stem peptides. These denuded glycan strands were then labeled at the reducing end by reductive amination with the anionic fluorophore 8-aminonaphthalene-1,3,6-trisulfonic acid (ANTS), and separated by gel electrophoresis for in-gel imaging (UV excitation at 365 nm, visible emission). **b**, In lane 1, showing PG isolated from a wild-type strain, shorter glycan strands are visualized as a discrete ladder. In lanes 2 and 3, representing glycan strands of Δ*sagB* and Δ*spdC* respectively, this discrete ladder of short glycan strands is lost.

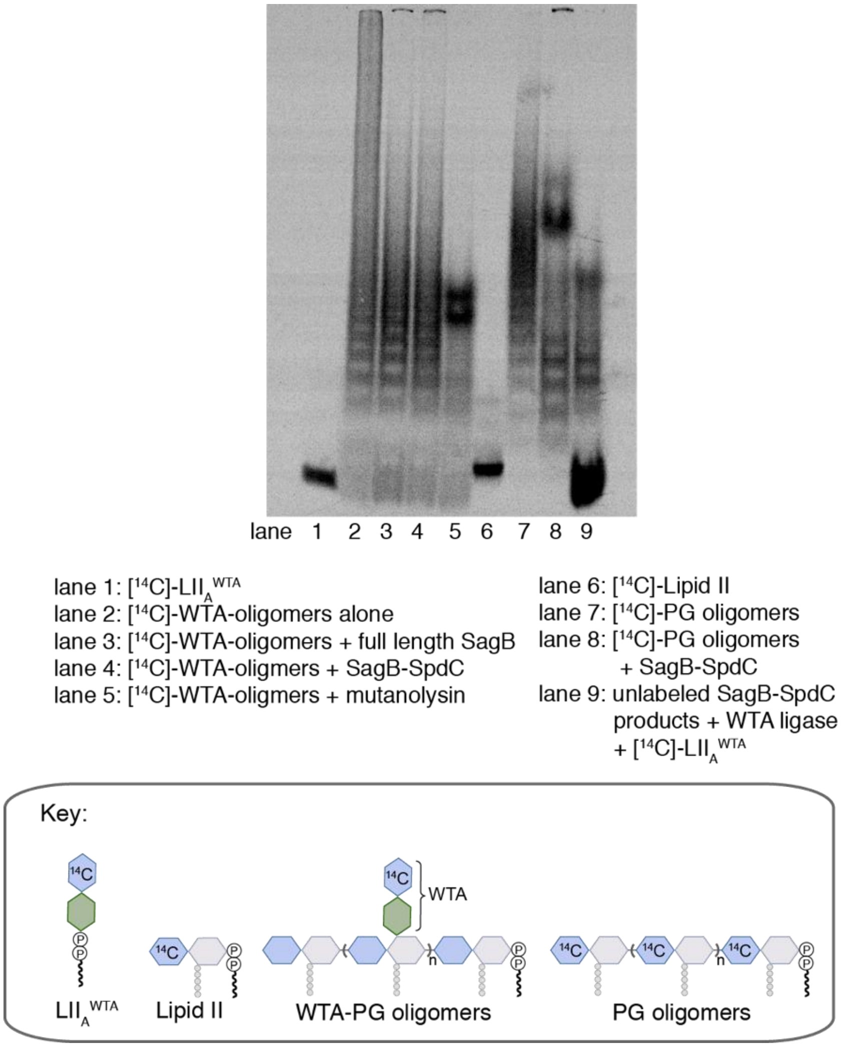


**Supplementary Figure 14. Nascent peptidoglycan is the preferred substrate of SagB-SpdC.** Cold glycan strands were prepared with radiolabeled wall teichoic acid (lanes 2-5) and incubated with a hydrolase. In the presence of either SagB (lane 3) or SagB-SpdC (lane 4), the ladder signal remained the same as a no treatment control (lane 2). There was reduced radiolabeled signal towards the top of the gel in both reactions; this signal represents a population of longer peptidoglycan oligomers and the fainter signal suggests less wall teichoic acid present. In contrast, mutanolysin was able to hydrolyze wall teichoic acid-modified glycan strands. To test whether SagB-SpdC cleavage interfered with wall teichoic acid ligation, cold oligomers were prepared and then incubated with radiolabeled wall teichoic precursor, $LII_A^{WTA}$, and the wall teichoic acid ligase, TagT. The appearance of the radiolabeled ladder indicates that SagB-SpdC products were ligated with wall teichoic acid (lane 9).

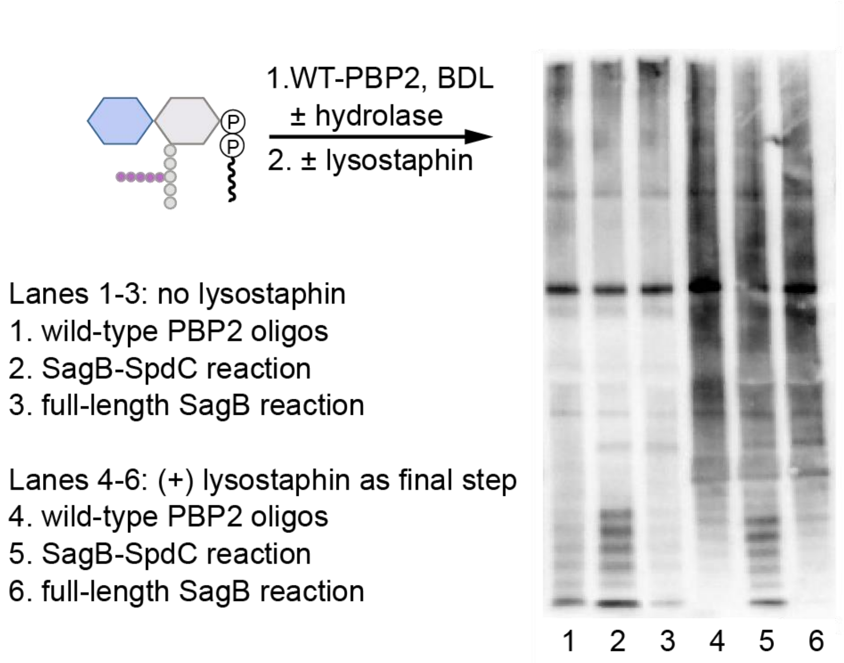


**Supplementary Figure 15. Crosslinked PG is not a substrate for SagB-SpdC.** *S. aureus* Lipid II was co-incubated with wild-type PBP2, a hydrolase, and BDL for visualization. Reactions were quenched, halved, and then one portion was treated with lysostaphin. Crosslinked peptidoglycan oligomers do not readily enter the gel matrix (lane 1) and appear as a dark smear when crosslinks are cleaved by the endopeptidase lysostaphin (lane 4). In reactions containing SagB-SpdC, a low-molecular weight ladder of bands is present without lysostaphin (lane 2) or with lysostaphin (lane 5). This shows that these products, generated by SagB-SpdC, are not crosslinked as they were not changed with lysostaphin treatment. Notably, approximately 20% of peptidoglycan peptides are crosslinked by wild-type PBP2 in vitro[11]. The high molecular weight signals (smears) in lanes 5 and 6 are similar to the signal in lane 4, indicating that crosslinked PG is not cleaved by either SagB-SpdC or full-length SagB.

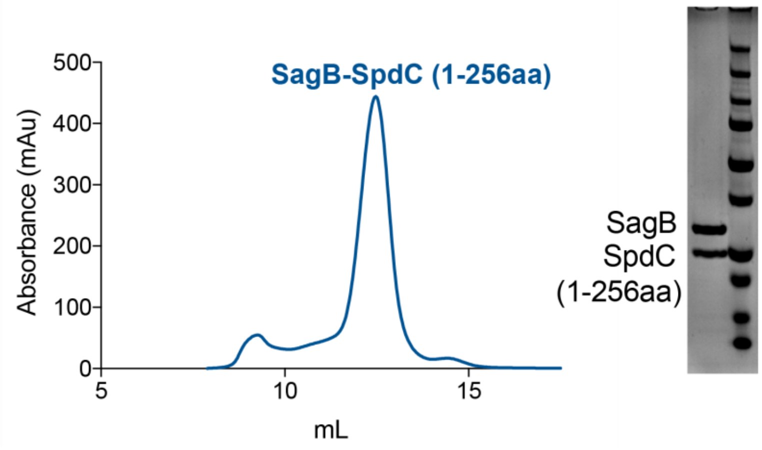


**Supplementary Figure 16. SagB and a truncated SpdC variant lacking its C-terminal, cytoplasmic region (SpdC$^{1\text{-}256}$) co-purify as a stable complex.** Expression and purification conditions used to obtain SagB-SpdC$^{1\text{-}256}$ complex were similar to those used in purification of the wild-type SagB-SpdC complex. A stable 1:1 protein complex was obtained after tandem affinity purifications and indicates that the C-terminal cytoplasmic region of SpdC is not required for complexation. *In vitro* and *in vivo* activity of this complex is also similar to that of wild-type SagB-SpdC (see Supplementary Fig. 11 and 17).

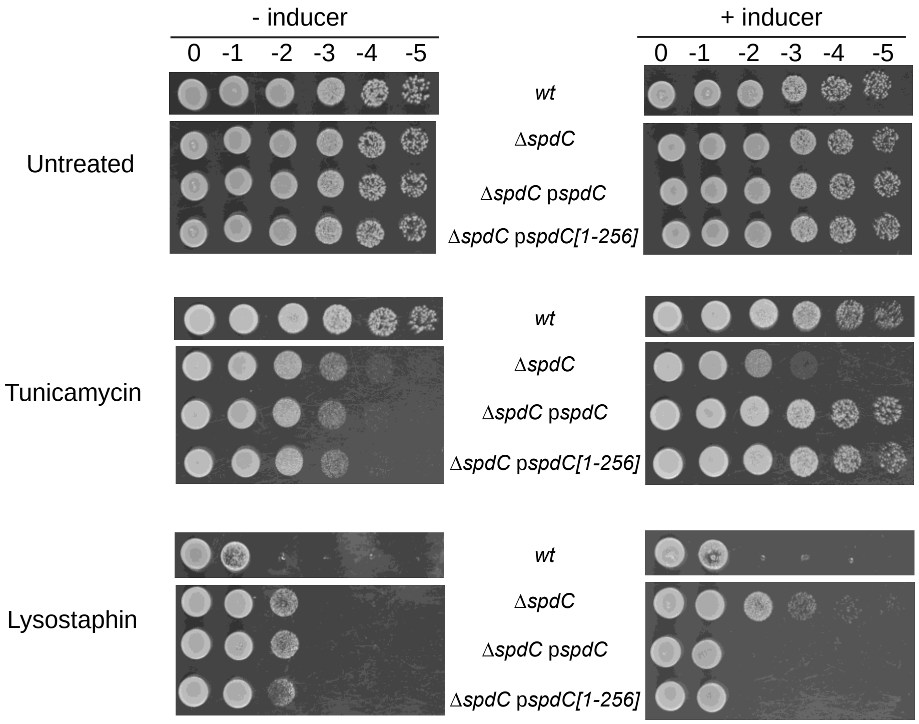


**Supplementary Figure 17. The cell wall phenotypes of SpdC are unaffected by its cytoplasmic truncation**. Spot dilution series performed as described in Supplementary Figure 12.

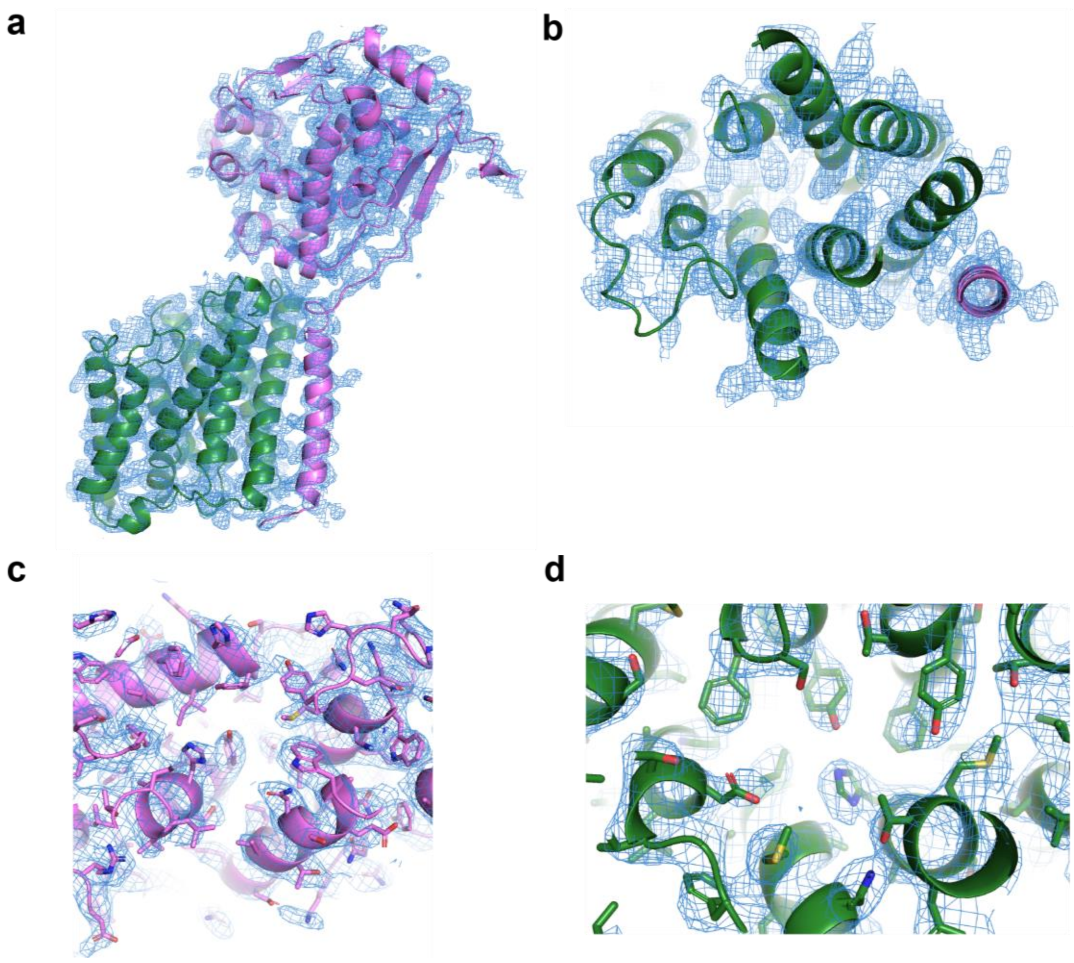


**Supplementary Figure 18. Representative electron density. a-d,** Simulated annealing composite omit $2F_O$-$F_C$ electron density maps, with SagB colored violet and SpdC colored green**. a,** Overall structure of SagB-SpdC viewed in the plane of the membrane. **b,** View of the transmembrane helices from the extracellular face. **c,** Active site groove of SagB, with sidechains shown as sticks. **d,** Central region of SpdC between several transmembrane helices in the same orientation as shown in b.

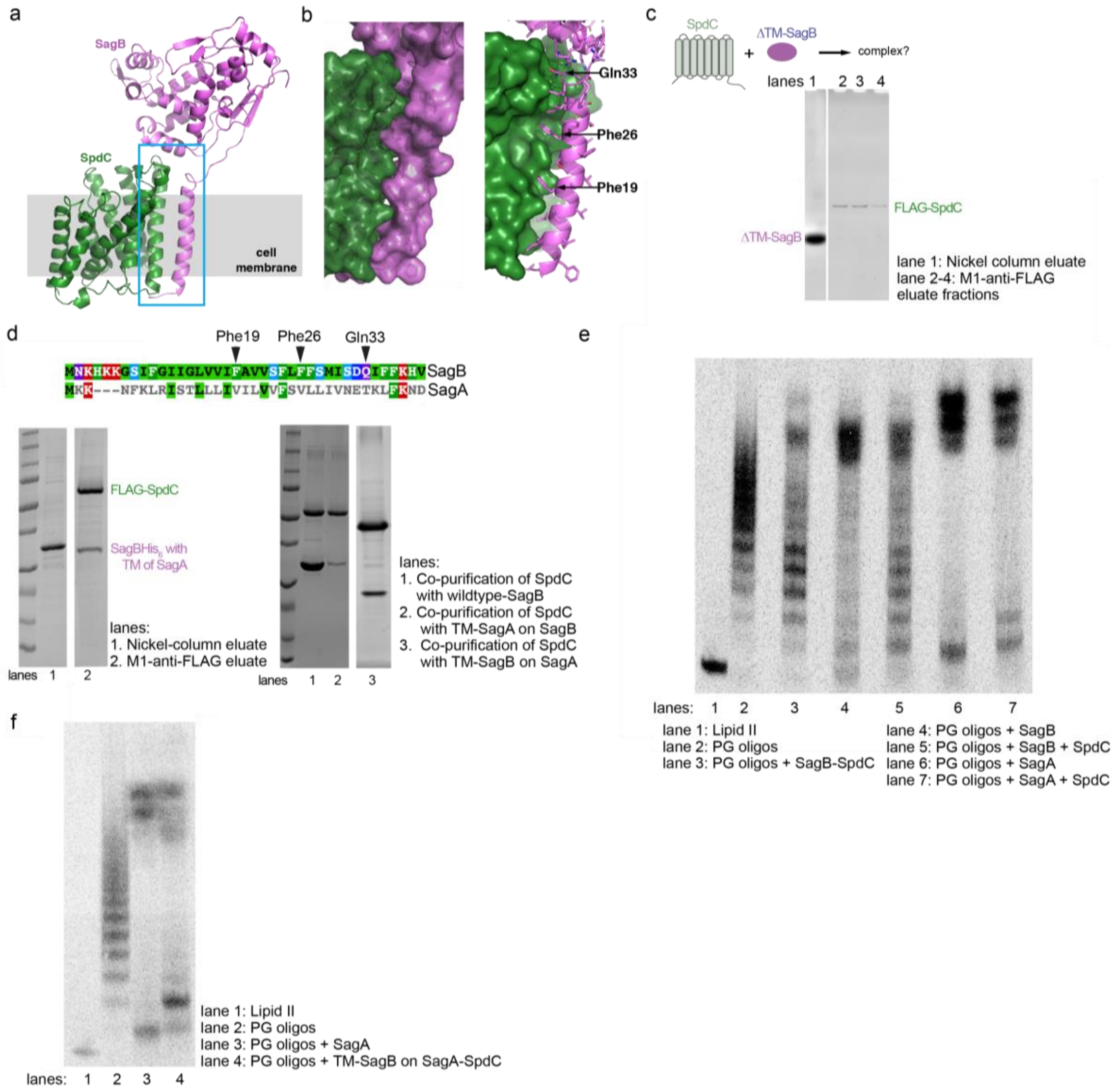


**Supplementary Figure 19. The interface between the transmembrane helices of SagB and SpdC is crucial for complexation. a,** Cartoon representation of the SagB-SpdC complex shows that the transmembrane (TM) helix of SagB is making close contacts to the TM3 of SpdC. **b,** Close-up of the interface between the SagB TM helix and SpdC. Left panel: Surface representation, Right panel: the same view showing the side-chains of the SagB TM helix as sticks. **c,** An attempt to co-purify ΔTM-SagB with SpdC did not yield a complex, although the individual components could be recovered after their corresponding affinity purification steps, as shown in SDS-PAGE. **d,** Alignment of sequences corresponding to the TM helices of SagA and SagB shows significant sequence differences. A construct of SagB containing the TM helix of SagA was co-expressed with SpdC. A stable complex did not form between SpdC and SagB with the TM of SagA, and very little of this SagB variant co-purified along SpdC as shown in SDS-PAGE. Inversely, a construct of SagA containing the TM of SagB was co-expressed with SpdC and did not form a stable 1:1 complex (right gel). **e,** Radiolabeled peptidoglycan oligomers treated with SagB-SpdC, individual hydrolases, or hydrolases with the addition of SpdC. Unlike SagB, SagA activity is not significantly altered by the presence of SpdC. **f,** Swapping the TM of SagB onto SagA has only a minor effect on its product distribution.

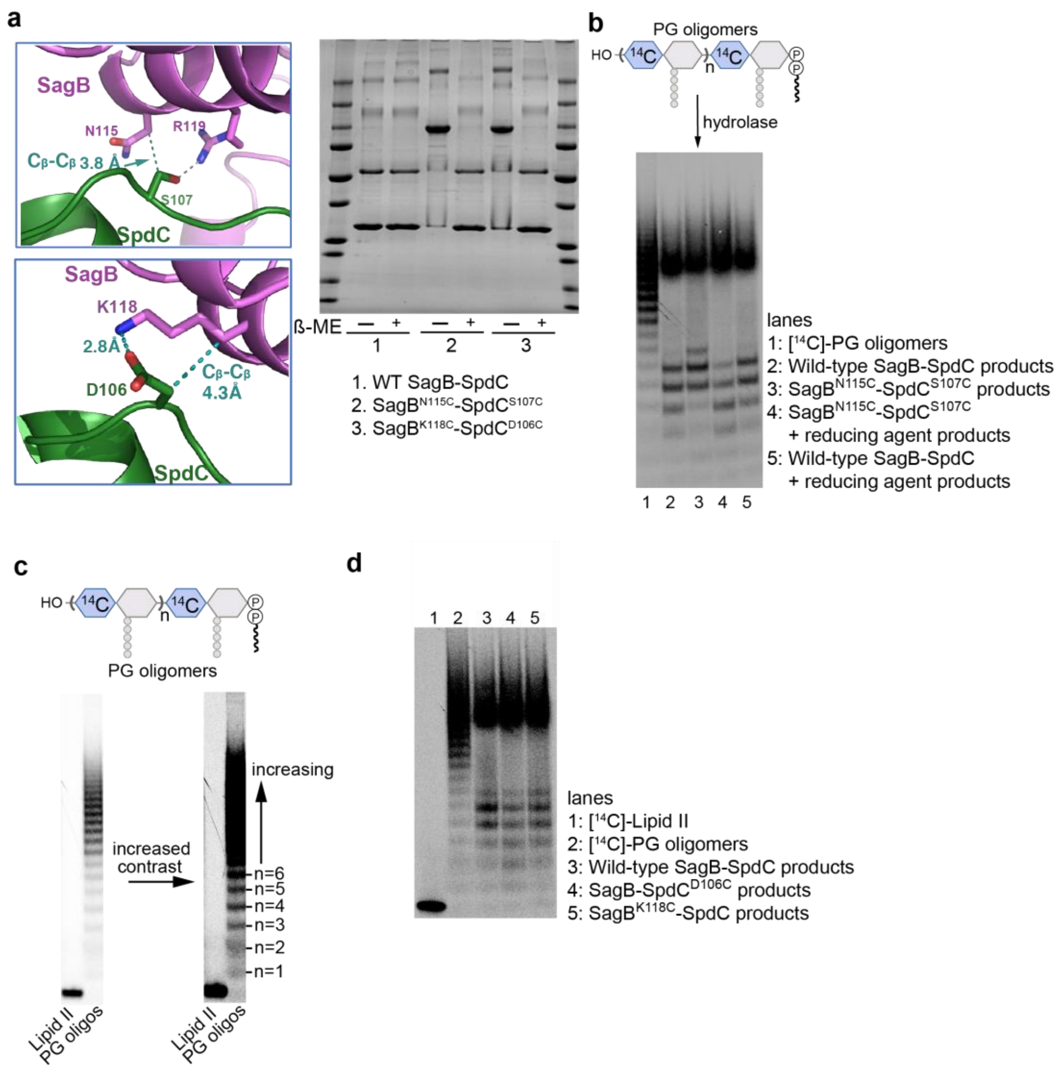


**Supplementary Figure 20. Disulfide-stabilized SagB-SpdC complexes cleave nascent peptidoglycan to a longer distribution of lipid-linked oligomers than the wild-type complex. a,** Close-up view of two interactions at the extracellular interface between SagB (violet) and SpdC (forest green). Sidechains of the relevant residues are shown as sticks. SpdC serine 107 hydrogen bonds to SagB arginine 119, however, its $C_\beta$ is closest to the $C_\beta$ of SagB asparagine 115. Wild-type SagB-SpdC, SagB$^{N115C}$-SpdC$^{S107C}$, and SagB$^{K118C}$-SpdC$^{D106C}$ were purified and then samples were mixed with 2x SDS-PAGE loading buffer with or without β-mercaptoethanol (β-ME), run on an acrylamide gel, and stained with coomassie. **b,** To test the effects of the cysteine substitutions and disulfide formation, radiolabeled PG oligomers (lane 1) were incubated with wild-type SagB-SpdC or SagB$^{N115C}$-SpdC$^{S107C}$ in the presence or absence of reducing agent. **c,** To assess the length (in sugar or disaccharide units) of lipid-linked, linear peptidoglycan fragments, we compare samples to Lipid II and the ladder of PG oligomers produced from it by PBP2. Each “step” in the ladder corresponds to the addition of a MurNAc-GlcNAc disaccharide. **d,** Single cysteine mutations in SagB-SpdC do not alter the distribution of cleavage products.

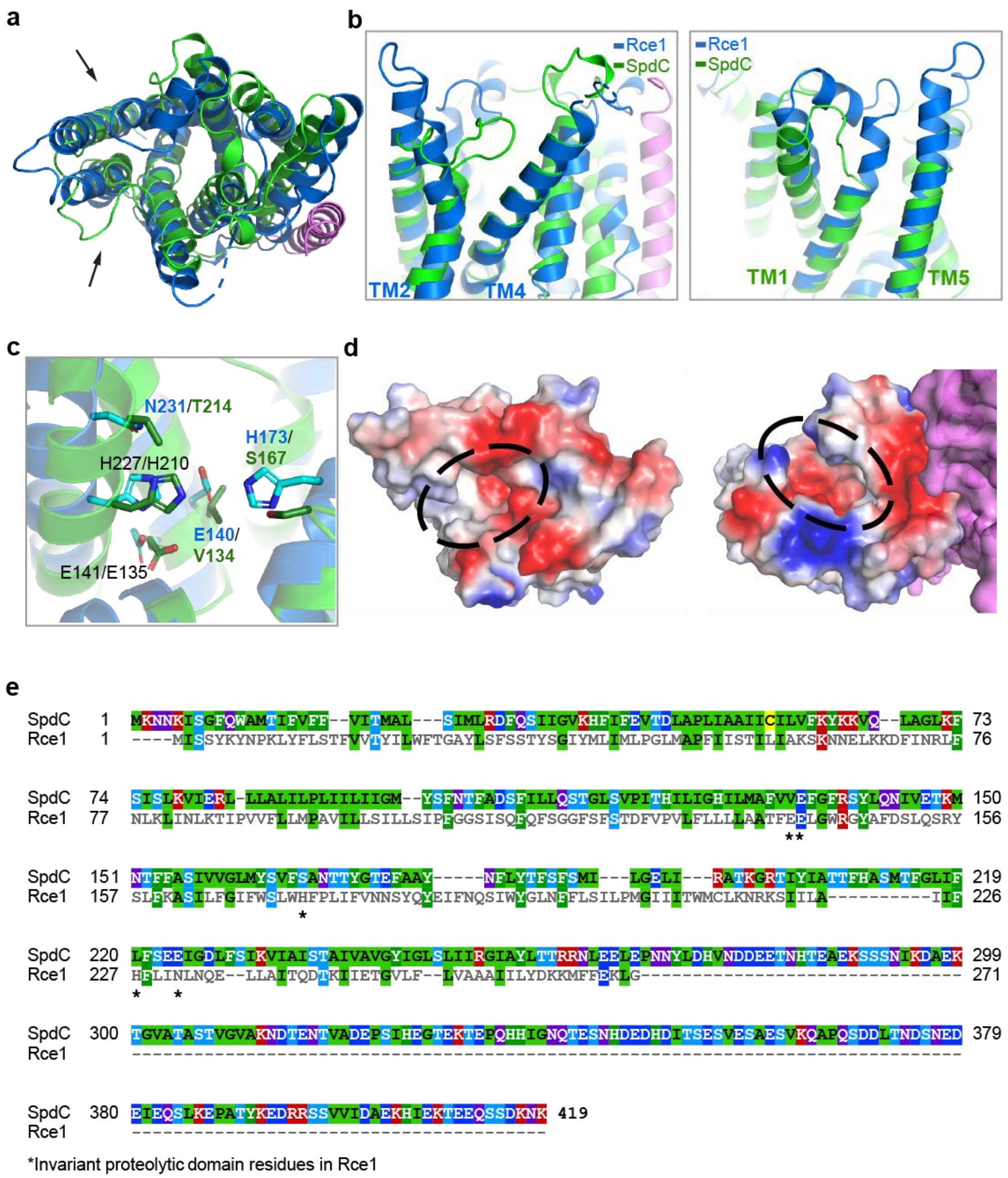


**Supplementary Figure 21. Alignment of SpdC and Rce1 structures shows broad similarities in their transmembrane domains, but key differences in the predicted substrate binding site. a**, Cartoon representation of the transmembrane helices of SagB-SpdC (violet, green) aligned with the structure of Rce1 (blue, PDB# 4CAD), with arrows denoting the views in b. **b**, The aligned structures when viewed from within the plane of the membrane. Although the helices of Rce1 and SpdC are threaded in the same pattern, the helices differ in length and the loops between helices adopt significantly different conformations. **c**, Several Rce1 catalytic residues (blue sticks) are not present in SpdC (green sticks); conserved residues are labeled in black. **d**, Surface electrostatic maps reveal that Rce1 (left) and SpdC (right) both contain deep central cavities that open to the membrane (circled in black). Both Rce1 and SagB-SpdC are oriented as in a, showing that the cavities open to the membrane at different locations and that the rim of the SpdC cavity contains more positively charged residues. **e**, Sequence alignment of SpdC and Rce1 generated using Clustal Omega and colored in MView. Stars under the alignment mark positions of the catalytic residues in Rce1.

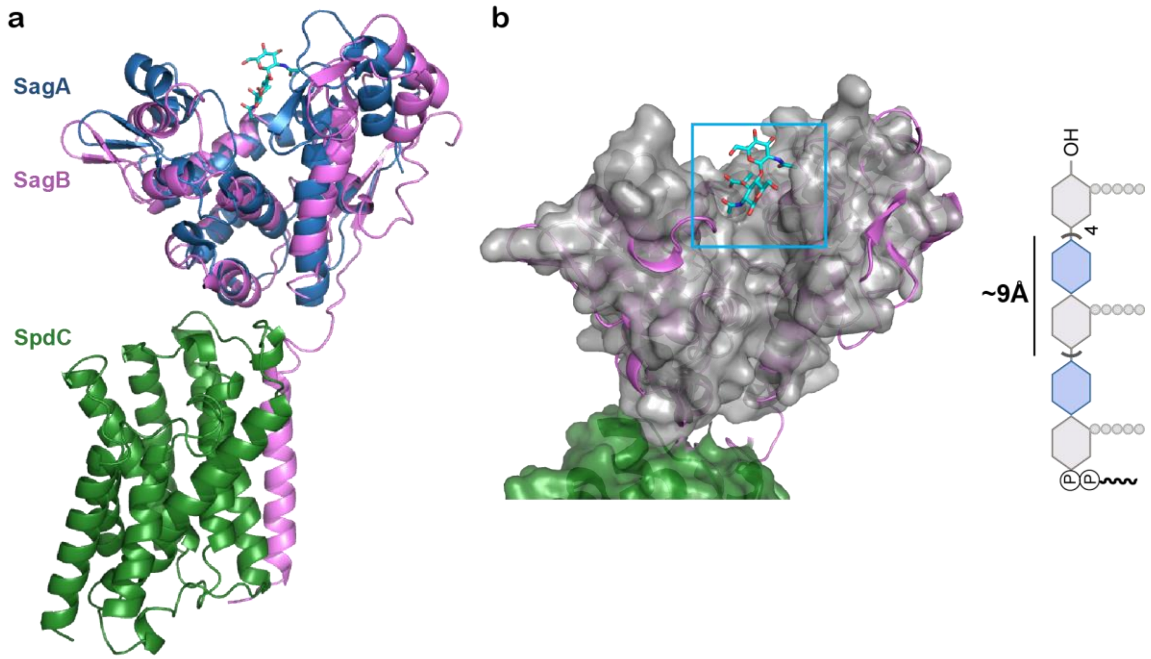


**Supplementary Figure 22. Alignment of SagB-SpdC with SagA crystal structures shows that the reducing end of a substrate glycan strand would be adjacent to the SagB-SpdC interface. a,** A crystal structure of SagA (blue) in complex with GlcNAc-MurNAc fragment (sticks) (PDB #4PI7) was aligned to SagB-SpdC (violet and forest green) using the glucosaminidase domain of SagB as the target. The orientation of the disaccharide fragment indicates that the reducing end faces SpdC and that the non-reducing end of the peptidoglycan substrate faces the top of the active site cleft. **b,** The same alignment shown in **a**, with SagA shown as a gray surface. The length of a disaccharide unit in the linear glycan chain is approximately 9 angstroms.

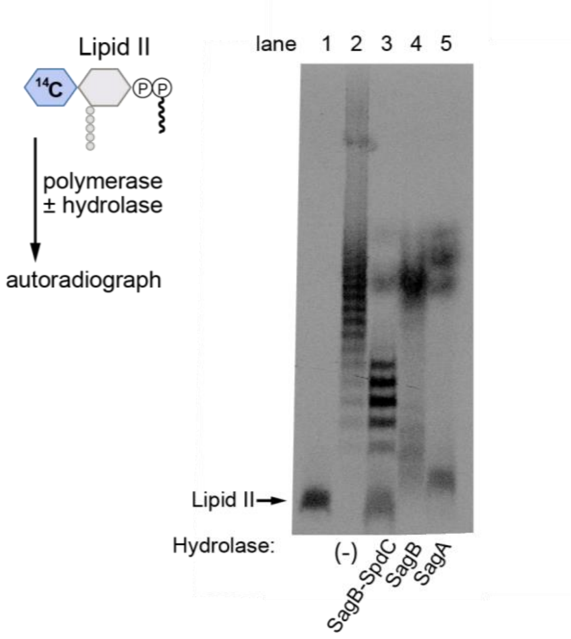


**Supplementary Figure 23. SagB-SpdC cleaves peptidoglycan oligomers with an active polymerase present.** Synthetic, [$^{14}$C]-Lipid II was incubated with PBP2 and a hydrolase (SagB-SpdC, SagB, or SagA). Similar cleavage products are observed in the presence of pre-assembled peptidoglycan oligomers (see main text Fig. 2b).

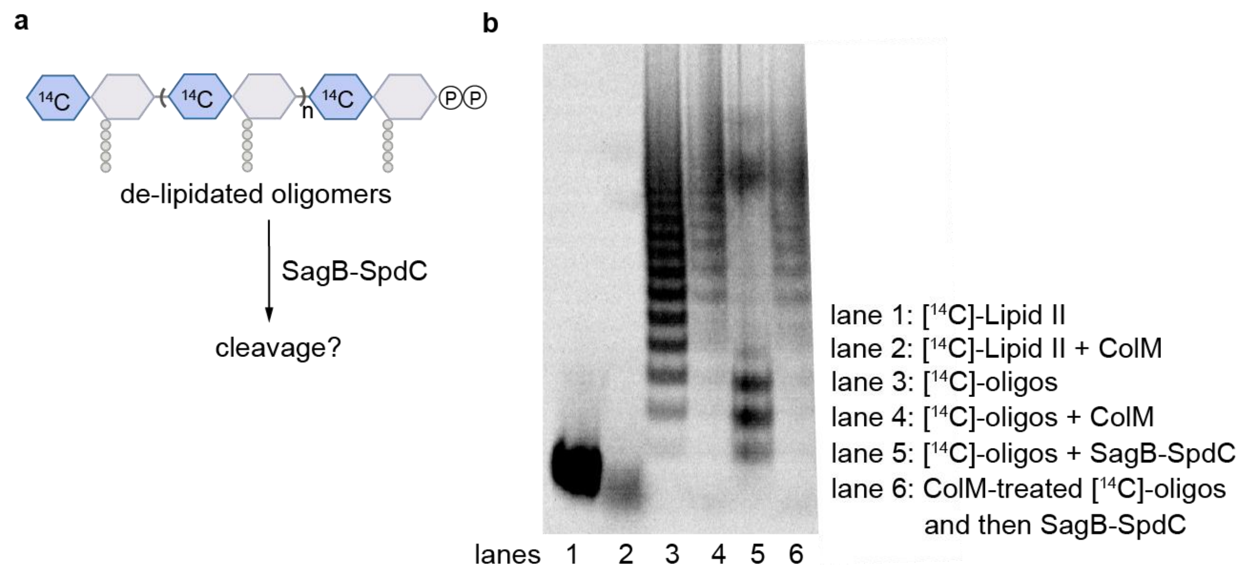


**Supplementary Figure 24. Removal of the lipid carrier from nascent peptidoglycan inhibits SagB-SpdC cleavage.** Radiolabeled peptidoglycan oligomers (lane 3) were treated with Colicin M (lane 4), which removes the lipid carrier, leaving a pyrophosphate on the reducing terminus of the glycan. Lipid-linked and de-lipidated oligomers were then treated with SagB-SpdC (lanes 5 and 6, respectively). Comparison of lane 3 to lane 5 shows changes to the sample characteristic of SagB-SpdC activity, whereas comparison of lane 4 to lane 6 shows no change, indicating that SagB-SpdC was not able to cleave oligomers lacking the lipid carrier.

| Data Collection | SagB-SpdC |
|---|---|
| Space Group | C2 |
| Unit Cell | |
| Dimensions (a, b, c), Å | 156.7, 54.8, 96.4 |
| Angles (α, β, γ), ° | 90.0, 107.2, 90.0 |
| Resolution range, Å | 46.98 - 2.60 (2.71 - 2.60) |
| Completeness, % | 99.3 (97.2) |
| $R_{merge}$ | 0.16 (1.86) |
| Mean I/σ(I) | 4.7 (0.8) |
| Multiplicity | 5.8 (5.1) |
| $CC_{1/2}$ | 0.994 (0.530) |
| **Refinement** | |
| No. unique reflections | 24207 |
| Rwork, % / Rfree, % | 25.3 / 27.9 |
| Average B (Å$^2$ ) protein | 98.7 |
| Average B (Å$^2$ ) ligands | 121.8 |
| Average B (Å$^2$ ) waters | 87.7 |
| Ramachandran plot | |
| Favored/disallowed (%) | 96.82 / 0.00 |
| rmsd from ideal geometry | |
| Bond lengths, Å | 0.0025 |
| Bond angles, ° | 0.49 |
| PDB code | 6U0O |

**Supplementary Table 3. Data collection and refinement statistics for SagB-SpdC$^{1\text{-}256}$.**

| Primer Name | Sequence (3'-5') |
|---|---|
| SM1 | GATCTGATTGGATCCAATATTGAGCATACGCAATATC |
| SM2 | CTAGCAAGCGCTTTGTATATGTAACCTCCATTAG |
| SM3 | CTAATGGAGGTTACATATACAAAGCGCTTGCTAG |
| SM4 | GATCTGATTGTCGACCATGACGCTGGGAATTGG |
| SM43 | GATCTGATTGGATCCCATAATGACATCATGTTCATGTAC |
| SM44 | CCTATCACCTCAAATGGTTCGCTATATGTAACCTCCATTAGG |
| SM45 | CCTAATGGAGGTTACATATAGCGAACCATTTGAGGTGATAGG |
| SM46 | GAGGTACTAGCAAGCGCTTTGTCCTAGGTACTAAAACAATTCATC |
| SM47 | GATGAATTGTTTTAGTACCTAGGACAAAGCGCTTGCTAGTACCTC |
| SM48 | GATCTGATTGTCGACGCAGAGCGCATCGGTCTTTTCG |
| SM124 | GATTGGTACCGTTACCTAATGGAGG |
| SM125 | TTAAGCTCAGCTTATTAATGGTGATGGTGATGGTGTTTGTTTTTATC TGAAGATTGTTC |
| SM130 | CTGATGGCGTTCGTAGTAGCATTCGGATTCCGTTCATAC |
| SM131 | GTATGAACGGAATCCGAATGCTACTACGAACGCCATCAG |
| SM132 | GTAGTAGAATTCGGATTCGCATCATACTTACAAAATATTG |
| SM133 | CAATATTTTGTAAGTATGATGCGAATCCGAATTCTACTAC |
| SM134 | TATATTGCAACGACATTCGCAGCTTCAATGACATTCGGA |
| SM135 | TCCGAATGTCATTGAAGCTGCGAATGTCGTTGCAATATA |
| SM165 | GATTGGTACCAATGGAGGTTACATATAATGGAACAAAAACTTATTTC TGAAGAAGATCTGAAGAACAATAAAATTTCTGGTTTTCAATG |
| SM166 | GATTGCTCAGCTTATTATTTGTTTTTATCTGAAGATTGTTCTTC |
| oJP21 | CGGCGCTCAGCTCAGTGGTGGTGGTGGTGGT |
| oJP22 | TGATGGTACCAACAATGACCTAAGAGGTGTGGATATGAATAAACAC AAGAAAGGTTCTATTTTTGG |
| oJP25 | TGATGGTACCAATGGAGGTTACATATAA |
| oJP26 | CGGCGCTCAGCTTATTATTTGTTTTTAT |
| oJP30 | TGATGGTACCAATGGAGGTTACATATAATGGAGCAAAAACTTATTTC TGAAGAAGATC |
| oJP31 | CGGCGCTCAGCTTATTATTTGTTTTTATCTGAAGATTGTTCCTCAG |
| F_pKTarO | GCTTATCAAAGTCGACCTGCAGGCATGCAAGCTTGGCGTAATC |
| R_pkTarO | CAACAATGACCTAAGAGGTGTGGATGGATCCCCGGGTACCG |

| | |
|---|---|
| F_1kb+_sagB | AAGATATATGTTCAATAAAATAACTTAGTTGGGAATGGGCTTCATTTCAG |
| R_1kb+_sagB | GATTACGCCAAGCTTGCATGCCTGCAGGTCGACTTTGATAAGC |
| F_1kb(-)_sagB | CGGTACCCGGGGATCCATCCACACCTCTTAGGTCATTGTTG |
| R_1kb(-)_sagB | ATTGCGAGATTTGGGTTGTTGAGCCATAGTCTTTCTCTTTGATTTAAAAG |
| F_tetR_sagB | TTAAATCAAAGAGAAAGACTATGGCTCAACAACCCAAATCTCGCAATTTG |
| R_tetR_sagB | AAGCCCATTCCCAACTAAGTTATTTTATTGAACATATATCTTACTTTATC |
| oJP32 | CCGGGGATCCCGAAGGTCATGTTCATTAAAGTATGTTG |
| oJP33 | AACTCAAATTGCGAGATTTGGGTTGATCCACACCTCTTAGGTCATTGTTG |
| oJP35 | GCAGGTCGACCATCAATAATGAATAATATGCCTACAGTTTTAATA |
| oJP49 | TTTGATAAGCTACGAGTTGTTTTT |
| oJP50 | ATCCACACCTCTTAGGTCATTGT |
| oJP51 | CTAAGAGGTGTGGATCAACCCAAATCTCGCAAT |
| oJP52 | TCGTAGCTTATCAAACTAAGTTATTTTATTGAACAT |
| oJP53 | GGATCCCCGGGTACCGAG |
| oJP54 | GGTACCCGGGGATCCCGAAGGTCATGTTCATTAAAGTATG |
| oJP79 | TAGAGAATAGGAACTTCATCCACACCTCTTAGGTCATTGTTG |
| oJP80 | GAAAGTATAGGAACTTCTTTGATAAGCTACGAGTTGTTTTTATGACTC |
| oJP81 | CGGCGCTCAGCTTATGCAATACCTCGGATAATTAAGCTTAAAC |
| oTD73 | TATTCTCTAGAAAGTATAGGAACTTCGCGAACCATTTGAGGTGATAGGTAAGATTAT |
| oTD74 | TACTTTCTAGAGAATAGGAACTTCTCCTAGGTACTAAAACAATTCATCCAGTAA |
| oTD145 | CAACCCAAATCTCGCAATTTGAGTTG |
| F_SpdC | CATCCACTGAATATATGAAGAACAATAAAATTTCTGGTTTTCAATG |
| R_SpdC | GCTCGAATTCGGATCCTTATTTGTTTTTATCTGAAGATTGTTCTT |
| F_DUET_FLAG_SpdC | CAGATAAAAACAAATAAGGATCCGAATTCGAGCTCGG |
| R_DUET_FLAG_SpdC | GAAATTTTATTGTTCTTCATATATTCAGTGGATGACCCCCCAG |
| F_SagB | GAGATATACATATGAATAAACACAAGAAAGGTTCTATTTTTGG |

| | |
|---|---|
| R_SagB | GCAAGCTTCTTATTCAAATGTTTACTGTCATCTTTATACAC |
| F_DUET_SagB | ACAGTAAACATTTGAATAAGAAGCTTGCGGCCG |
| R_DUET_SagB | TCTTGTGTTTATTCATATGTATATCTCCTTCTTATACTTAACTAATATAC |
| R_Trunc_SpdC | CCGAGCTCGAATTCGGATCCTTATGCAATACCTCGGAT |
| F_DUET_trunc_SpdC | ATCCGAGGTATTGCATAAGGATCCGAATTCGAGCTCGG |
| F_SagB_alone | TTAAGAAGGAGATATACCATGAATAAACACAAGAAAGGTTCTATTTTTGG |
| R_SagA_alone | GCAAGCTTCTTATTCAAATGTTTACTGTCATCTTTATACAC |
| F_plasmid_SagA | ACAGTAAACATTTGAATAAGAAGCTTGCGGCCG |
| R_plasmid_SagA | AGAACCTTTCTTGTGTTTATTCATGGTATATCTCCTTCTTAAAGTTAAAC |
| F_soluble_SagB | ATGGCTAGCCAGATATTTTTCAAACATGTTAAATCCG |
| R_soluble_SagB | CATTTGCTGTCCACCAGTCATTTACTTATTCAAATGTTTACTGTCATC |
| F_plasmid_soluble_SagB | GATGACAGTAAACATTTGAATAAGTAAATGACTGGTGGACAGCAAATG |
| R_plasmid_soluble_SagB | GCTAGCCATATGGCTGCCG |
| F_SagB_NRIKRMLVD_to_SEVNQLLKG | CAAGGGATTGATAAA TCTGAAGTTAACCAATTGCTAAAAGGTAGACCAACGTTATTGAAACATACGGATG |
| R_SagB_NRIKRMLVD_to_SEVNQLLKG | GTTGGTCTACCTTTTAGCAATTGGTTAACTTCAGATTTATCAATCCCTTGATACTTTGATAAATCTAAAA |
| F_SagB_interface_mut | TCTAGAATTAATCAAATGTTAGACGGTAGACCAACGGAATTGAAACATACGGATGATTTCTTAAAAGCTG |
| R_SagB_interface_mut | TTC CGT TGG TCT ACC GTC TAA CAT TTG ATT AAT TCTAGATTTATC AAT CCC TTG ATA CTT TGA TAA ATC |
| F_SpdC_D106R | GGTATGTACAGCTTTAATACATTTGCAAGAAGCTTTATTTTATTACAATCAACAGGC |
| R_SpdC_D106R | GCC TGT TGA TTG TAA TAA AAT AAA GCT TCT TGC AAA TGT ATT AAA GCT GTA CAT ACC |
| F_SagB_K118D | GTATCAAGGGATTGATAAAAATAGAATTGATCGTATGTTAGTAGATAGACCAACG |
| R_SagB_K118D | CGTTGGTCTATCTACTAACATACGATCAATTCTATTTTTATCAATCCTTGATAC |
| R_pETduet_s2_SagB1-5 | TTTCTTGTGTTTATTCATATGTATATCTCCTTCTTATACTTAAC |
| F_pETduet_SagA5 | GTTAAGTATAAGAAGGAGATATACATATGAATAAACAC AAGAAAAATTTCAAGTTACGCATTTCAACGC |
| F_SagA_TM_with_SagB_overlap | CGGATTTAACATGTTTGAACAATTTAGTTTCATTCACGATGAGTAATACAGC |

| | |
|---|---|
| R_SagA_TM_with_SagB_overlap | CGGATTTAACATGTTTGAACAATTTAGTTTCATTCACGATGAGTAATACAGC |
| F_SagB_TM_then_SagB_F36 | CTCATTTTTATTTTTCTCAATGATATCCGATACTAAATTGTTTAAAAATGATGTGAATTACTC |
| R_SagB_TM_then_SagB_F36 | GAGTAATTCACATCATTTTTAAACAATTTAGTATCGGATATCATTGAGAAAAATAAAAATGAG |
| R_SpdC_D106C | GCCTGTTGATTGTAATAAAATAAAGCTGCATGCAAATGTATTAAAGCTGTACATACC |
| F_SpdC_S107C | GTACAGCTTTAATACATTTGCAGATTGCTTTATTTTATTACAATCAACAGGC |
| R_SpdC_S107C | GCCTGTTGATTGTAATAAAATAAAGCAATCTGCAAATGTATTAAAGCTGTAC |
| F_SagB_N115C | TTATCAAAGTATCAAGGGATTGATAAATGTAGAATTAAACGTATGTTAGTAGATAGAC |
| R_SagB_N115C | GTCTATCTACTAACATACGTTTAATTCTACATTTATCAATCCCTTGATACTTTGATAA |
| F_SagB_K118C | GTATCAAGGGATTGATAAAAATAGAATTTGCCGTATGTTAGTAGATAGACCAACG |
| R_SagB_K118C | CGTTGGTCTATCTACTAACATACGGCAAATTCTATTTTTATCAATCCC TTGATAC |

**Supplementary Table 4.  Primers used in this study**

| **Strain** | **Genotype** | **Source** |
|---|---|---|
| RN4220 *S. aureus* | Wild-type | [12] |
| TD011 | RN4220 *S. aureus* (pTP44) | [13] |
| HG003 *S. aureus* | Wild-type | [14] |
| TM283 | *S. aureus* USA300 cured of pUSA300HOUMR; Tn library host | [15] |
| SHM002 | HG003 Δ*spdC* | This study |
| SHM056 | HG003 Δ*spdC*::*kan*$^{R}$ | This study |
| SHM226 | HG003 Δ*spdC*::*kan*$^{R}$, pSM_*spdC*_Myc | This study |
| JP012 | HG003 *sagB::Tn-erm*$^{R}$ | This study |
| JP051 | HG003 *sagB::Tn-erm*$^{R}$ (pJP15) | This study |
| JP053 | HG003 *sagB::Tn-erm*$^{R}$ (pJP19) | This study |
| JP054 | HG003 Δ*spdC::kan*$^{R}$ (pJP17) | This study |
| JP061 | HG003 Δ*spdC*::*kan*$^{R}$ (pSM_*spdC*_E135A) | This study |
| JP062 | HG003 Δ*spdC*::*kan*$^{R}$ (pSM_*spdC*_R139A) | This study |
| JP063 | HG003 Δ*spdC*::*kan*$^{R}$ (pSM_*spdC*_H210A) | This study |
| JP064 | HG003 Δ*spdC*::*kan*$^{R}$ (pJP22) | This study |
| JP065 | HG003 Δ*spdC*::*kan*$^{R}$ *sagB::Tn-erm*$^{R}$ | This study |
| JP128 | HG003 Δ*spdC::kan*$^{R}$ (pJP42) | This study |
| JP132 | HG003 Δ*sagB::kan*$^{R}$ | This study |
| BL21(DE3) *E. coli* | *F- ompT hsdSB*(rB- mB-) *gal dcm* (DE3) | Novagen |
| NovaBlue(DE3) *E. coli* | *endA1 hsdR17* (rk12- mk12+) *supE44thi-1 recA1 gyrA96 relA1 lac* (DE3) *F' proAB lacI*$^{Q}$*ZΔM15::Tn10*(Tet$^{R}$)*gyrA96*(nal$^{R}$) *thi-1 recA1 relA1 lac glnV44 F'*[*::Tn10 proAB*$^{+}$ *lacI*$^{q}$ Δ(*lacZ*)*M15*] *hsdR17*(*rK– mK*+) | Novagen |
| **Plasmid** | **Description** | **Source** |

| | | |
|---|---|---|
| pTarKO | *cam*$^R$ vector for making gene deletions in *S. aureus* | [16] |
| pKFC | *cam*$^R$ vector for making gene deletions in *S. aureus* | [17] |
| pKFC_*spdC_kan* | pKFC vector to make Δ*spdC*::*kan*$^R$ marked deletion in *S. aureus* | This study |
| pKFC_*spdC* | pKFC vector to make Δ*spdC* unmarked deletion in *S. aureus* | This study |
| pTP63 | atc-inducible, integrative *cam*$^R$ expression vector for *S. aureus* | [13] |
| pSM_*spdC_myc* | pTP63-cMyc-*spdC* containing native *spdC* ribosome-binding site | This study |
| pSM_*spdC_his* | pTP63-*spdC*-His$_6$ containing native *spdC* ribosome-binding site | This study |
| pSM_*spdC*_E135A | pTP63-*spdC E135A*-His$_6$ containing native *spdC* ribosome-binding site | This study |
| pSM_*spdC* _R139A | pTP63-*spdC R139A*-His$_6$ containing native *spdC* ribosome-binding site | This study |
| pSM_*spdC* _H210A | pTP63-*spdC H210A*-His$_6$ containing native *spdC* ribosome-binding site | This study |
| pJP15 | pTP63-*sagB*-His$_6$ containing native *sagB* ribosome-binding site and 7 amino acid linker | This study |
| pJP17 | pTP63-cMyc-*spdC* containing native *spdC* ribosome-binding site | This study |
| pJP19 | pTP63-*sagB* E155A-His$_6$ containing native *sagB* ribosome-binding site and 7 amino acid linker | This study |
| pJP22 | pTP63-cMyc-*spdC* E135A/R139A/H210A containing native *spdC* ribosome-binding site | This study |
| pJP42 | pTP63-cMyc-*spdC[aa1-256]* containing native *spdC* ribosome-binding site | This study |
| pJP47 | pTarKO vector to make Δ*sagB::kan*$^R$ marked deletion in *S. aureus* | This study |
| pETDuet-FLAG | *amp*$^R$ vector for dual protein expression | [18] |
| pAM174 | *cam*$^R$ vector for Ulp1 protease expression | [19] |
| pET24b(+)-*sgtBY181D* | *amp*$^R$-marked *S. aureus* SgtB$^{WT}$-His$_6$ expression vector | [3] |
| pMW1010 | pET28b(+)-His6-*EF_3129* [T36-P429] expression vector for *E. faecalis* PBPX, *kan*$^R$ | [6] |
| p*pbp2*$^{S398G}$ | pET_42a containing a PBP2 construct with a S398G, TP inactive mutation | [8] |
| p*spdC_sagB* | pDUET containing a SUMO-FLAG-SpdC and a SagB-His$_6$ | This study |
| p*spdC_sagB with TM of SagA* | pDUET containing a SUMO-FLAG-SpdC and a SagB-His$_6$ with the TM of SagA | This study |
| p*spdC_sagA with TM of SagB* | pDUET containing a SUMO-FLAG-SpdC and a SagA-His$_6$ with the TM of SagB | This study |
| pspdC alone | pDUET containing only SUMO-FLAG-SpdC | This study |
| p*spdC*$^{E135A}$*_sagB* | pDUET containing a SUMO-FLAG-SpdC with E135A mutation and a SagB-His$_6$ | This study |
| p*spdC_sagB*$^{E155A}$ | pDUET containing a SUMO-FLAG-SpdC and a SagB-His$_6$ with a E155A mutation | This study |

| | | |
|---|---|---|
| pDUET_*spdC*$^{D106R}$_*sagB* | pDUET containing a SUMO-FLAG-SpdC with a D106R mutation and a SagB-His$_6$ | This study |
| pDUET_*spdC_sagB* *interface* mutant* | pDUET containing a SUMO-FLAG-SpdC and a SagB-His$_6$ the following mutations: N115S, K118N, R119Q, V122D, D123G, L127E | This study |
| p*spdC*$^{S107C}$_*sagB*$^{N115C}$ | pDUET containing a cysteine pair SUMO-FLAG-SpdC (S107C) and a SagB-His$_6$ with a N115C | This study |
| p*spdC*$^{D106C}$_*sagB*$^{K118C}$ | pDUET containing a cysteine pair SUMO-FLAG-SpdC (D106C) and a SagB-His$_6$ with a K118C | This study |
| p*sagB* | pDUET containing a full-length SagB-His$_6$ | This study |
| p*sagB*$^{(33\text{-}end)}$ | pET_28(b-) containing a soluble, SagB construct lacking the transmembrane domain (33-end aa) | This study |
| p*spdC* | pDUET containing a SpdC with an amino terminal sumo and flag epitope fusion | This study |
| p*spdC*$^{truncated}$_*sagB* | pDUET containing a truncated SpdC lacking the cytoplasmic carboxy terminus (1-253 aa) with an amino terminal Sumo and flag epitope fusion and a SagB-His$_6$ | This study |
| p*spdC_sagA* | pDUET containing a SpdC with an amino terminal sumo and flag epitope fusion and a SagA-His$_6$ | This study |

**Supplementary Table 5. Strains and plasmids used in this study**
